# Systematic annotation of orphan RNAs reveals blood-accessible molecular barcodes of cancer identity and cancer-emergent oncogenic drivers

**DOI:** 10.1101/2024.03.19.585748

**Authors:** Jeffrey Wang, Jung Min Suh, Brian J Woo, Albertas Navickas, Kristle Garcia, Keyi Yin, Lisa Fish, Taylor Cavazos, Benjamin Hänisch, Daniel Markett, Shaorong Yu, Gillian Hirst, Lamorna Brown-Swigart, Laura J. Esserman, Laura J. van ‘t Veer, Hani Goodarzi

**Affiliations:** Department of Biochemistry & Biophysics, University of California, San Francisco, San Francisco, California, USA; Department of Urology, University of California, San Francisco, San Francisco, California, USA; Helen Diller Family Comprehensive Cancer Center, University of California, San Francisco, San Francisco, California, USA; Bakar Computational Health Sciences Institute, University of California, San Francisco, CA, US Department of Surgery, University of California, San Francisco, San Francisco, CA 94143, USA; Biological and Medical Informatics, University of California San Francisco, San Francisco, CA, 94158, USA; Department of Surgery, University of California, San Francisco, San Francisco, CA 94143, USA; Department of Laboratory Medicine, University of California, San Francisco, San Francisco, CA 94143, USA; Arc Institute, Palo Alto, CA 94304, USA; School of Medicine, University of California, Davis, CA, US; Institut Curie, CNRS UMR3348, INSERM U1278, Orsay, France

## Abstract

From extrachromosomal DNA to neo-peptides, the broad reprogramming of the cancer genome leads to the emergence of molecules that are specific to the cancer state. We recently described orphan non-coding RNAs (oncRNAs) as a class of cancer-specific small RNAs with the potential to play functional roles in breast cancer progression^1^. Here, we report a systematic and comprehensive search to identify, annotate, and characterize cancer-emergent oncRNAs across 32 tumor types. We also leverage large-scale *in vivo* genetic screens in xenografted mice to functionally identify driver oncRNAs in multiple tumor types. We have not only discovered a large repertoire of oncRNAs, but also found that their presence and absence represent a digital molecular barcode that faithfully captures the types and subtypes of cancer. Importantly, we discovered that this molecular barcode is partially accessible from the cell-free space as some oncRNAs are secreted by cancer cells. In a large retrospective study across 192 breast cancer patients, we showed that oncRNAs can be reliably detected in the blood and that changes in the cell-free oncRNA burden captures both short-term and long-term clinical outcomes upon completion of a neoadjuvant chemotherapy regimen. Together, our findings establish oncRNAs as an emergent class of cancer-specific non-coding RNAs with potential roles in tumor progression and clinical utility in liquid biopsies and disease monitoring.

## Introduction

Cancer-emergent macromolecules, defined as molecules that are uniquely present in cancer cells, have become the focus of many studies in recent years. Structural variations that lead to the expression of cancer-specific fusion proteins have long been known to play a major role in tumorigenesis^2–4^. Tumors have also been shown to generate neoantigens, cancer-specific peptides that are absent in normal tissue, through the disruption of various cellular mechanisms^5,6^.

Extrachromosomal DNA (ecDNA) is another class of cancer-emergent molecules that can drive oncogenesis^7,8^. We previously reported the discovery of orphan non-coding RNAs (oncRNAs) in breast cancer, small non-coding RNAs that are expressed in cancer cells but are absent in non-transformed tissue^1^. We showed that one oncRNA, a small RNA derived from the TERC transcript, plays a functional role in breast cancer metastasis by disrupting a miRNA-mRNA regulatory network controlling the expression of prometastatic genes^1^. However, the extent to which oncRNAs may contribute functional roles in tumor progression across tumor types remains largely unexplored. In this study, we set out to systematically annotate oncRNAs across human cancers and discovered a large set of oncRNAs that are not only cancer-emergent but also cancer-specific and therefore provide a digital molecular barcode that can reliably discriminate different cancer types or even subtypes. Furthermore, we developed a large-scale *in vivo* genetic screening strategy to identify driver oncRNAs in multiple xenograft models of cancer. We discovered and subsequently validated several functional oncRNAs that impact tumor growth, indicating that they may have roles in disease progression.

We had previously shown that a fraction of oncRNAs are actively secreted by breast cancer cells and can potentially serve as a cancer-specific signal to distinguish serum samples from breast cancer and healthy patients. However, whether this signal was sufficiently strong to inform clinical practice in minimally-invasive clinical applications was unknown. Here, we found that many of the newly annotated oncRNAs are also actively secreted across different cancers, implying that this oncRNA molecular barcode is partially blood-accessible and can provide an opportunity for a sensitive and versatile liquid biopsy strategy for multiple cancers. In a first-in-class application of oncRNAs to liquid biopsy in minimal residual disease (MRD) detection, we performed a large retrospective analysis of breast cancer patients in an neoadjuvant chemotherapy setting. We demonstrated that cell-free oncRNAs provide a tumor-naive strategy for MRD applications in breast cancer with minimal sample volume and limited depth of sequencing. Altogether, our study encapsulates the first comprehensive effort to annotate oncRNAs across human cancers and reveal their potential as digital biomarkers for cancer cell identity, functional macromolecules in cancer progression, and blood-accessible, prognostic biomarkers. This work sets the stage for future investigations into the roles of oncRNAs in cancer biology and their applications in precision oncology strategies.

## Results

### Systematic annotation of orphan non-coding RNAs across human cancers

To systematically discover and annotate orphan non-coding RNAs, we started with raw small RNA sequencing data from the full The Cancer Genome Atlas (TCGA) dataset, which consists of roughly 10,400 tumor biopsies across 32 cancer types and 679 tumor-adjacent normal samples across 23 tissue types^9^. We first generated read-clusters by merging overlapping reads across all samples. We defined oncRNAs as those read-clusters that are significantly detected among the samples of a given cancer but are largely absent from the normal samples across all tissues. Because TCGA lacks data from most blood cancers and non-cancerous biofluids, we first used smRNA sequencing data from non-cancerous samples in the Extracellular RNA Atlas (exRNA Atlas) to filter our read-clusters (**Fig S1A**)^10^. We then removed read-clusters present in more than 10% of the TCGA tumor-adjacent normal samples for any of the tissue types. We systematically assessed the cancer-specific expression of the remaining smRNAs by using Fisher’s exact test to compare cancer samples from each tissue type against tumor-adjacent normal samples from all tissue types. Loci that were significant after multiple testing correction in at least one cancer type were annotated as oncRNAs.

By applying this framework, we discovered roughly 260,000 high-confidence oncRNA loci that are specifically expressed in one or more cancers (**Fig 1A** and **Fig S1B**). For example, we annotated 15,827 oncRNAs in breast cancer (TCGA-BRCA) and analyzed their presence and expression across both breast cancer and tumor-adjacent normal samples across all tissue types (**Fig S1C–D**). Overall, we annotated between 10^4^ and 10^5^ oncRNA species for each cancer type in TCGA (**Fig S1E**); some oncRNAs were unique to specific cancers while others were detected in more than one cancer (**Fig 1B**). Despite the low prevalence of any single oncRNA across all cancer samples (**Fig S1F)**, we observed that the binary patterns of presence and absence of multiple oncRNAs, which we have named oncRNA fingerprints, are readily distinguishable between cancer types. Comparing the median Jaccard similarity of oncRNA fingerprints between samples from the same cancer tissue type versus all other cancer tissue types, we found significantly higher similarity among samples from the same tissue-of-origin (**Fig S1G).** Therefore, each cancer type can be represented as a barcode based on the pattern of expressed oncRNAs (**Fig 1A, C** and **Fig S1H**).

**Figure 1.**
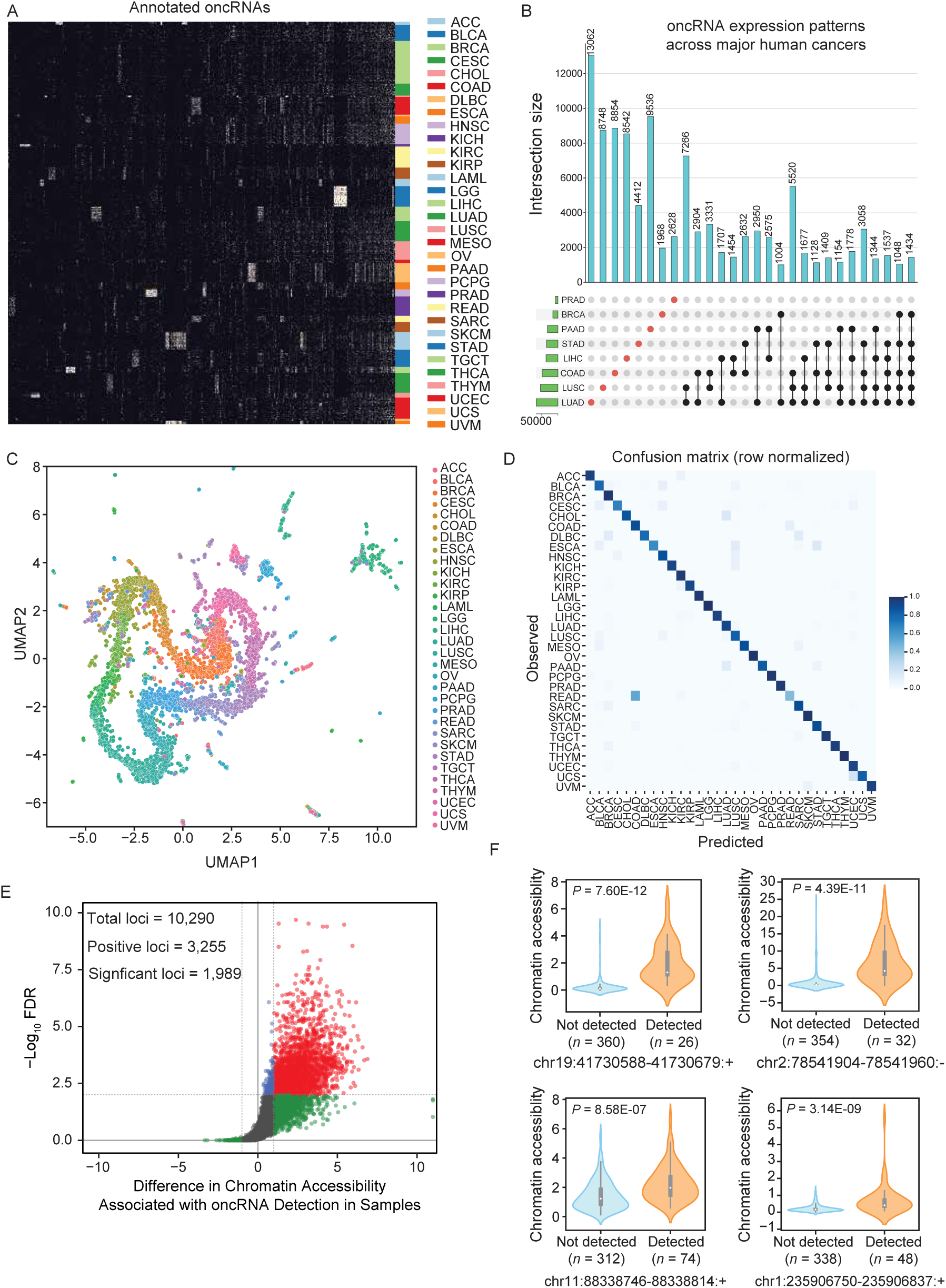
Systematic annotation of oncRNA loci across human cancers using small RNA sequencing data from TCGA and exRNA atlas. **(A)** A binary heatmap representing the presence and absence of oncRNA species across human cancers. Here we show a subset of 2,808 of the top significant oncRNAs. The subset was created by selecting 100 of the most significant oncRNAs for each cancer type as determined by the Fisher exact test and collapsing oncRNAs selected multiple times. Each column represents an annotated oncRNA, and each row represents one TCGA sample. Rows were grouped based on their tumor type (TCGA code) and columns were clustered based on their patterns. **(B)** Number of oncRNAs associated with the major human cancers, namely lung, breast, and gastrointestinal cancers, depicted as an UpSet plot. The vertical blue bars represent the oncRNA counts across one or more cancers with the exact numbers included at the top. **(C)** A 2D UMAP projection summarizing the oncRNA profiles across TCGA cancer samples. Samples are colored by tumor type. **(D)** The confusion matrix for tissue-of-origin classification based on oncRNA presence and absence in each sample. The matrix was row-normalized. **(E)** A volcano plot representing the relationship between chromatin accessibility and oncRNA detection. The x-axis represents, for each oncRNA, the log2 median difference in chromatin accessibility between samples in which the oncRNA was present versus absent. The y-axis shows the significance of the observed differences based on FDR corrected *P* values calculated using a one-sided Mann-Whitney test. A total of 10,290 oncRNA loci were considered for this analysis based on the coverage of ATAC data. Of these, 3,255 showed a positive association between oncRNA presence and increased chromatin accessibility; of these, 1,989 were also statistically significant at an FDR of 1%. **(F)** Chromatin accessibility signal of four exemplary oncRNA loci from (E), grouped by the detection of the cognate oncRNA in the small RNA dataset of each sample. Values are shown as violin plots and boxplots. The boxplots show the distribution quartiles, and the whiskers show the quartiles ± IQR (interquartile range). Also reported are the number of samples in which the oncRNAs were detected as well as their associated corrected *P* values.

To formalize this relationship, we took advantage of machine learning-based classifiers to assess the extent to which the oncRNA fingerprint from a given sample could be used to identify its tissue-of-origin (TOO). For this task, we first split the samples from TCGA into train and test datasets (80:20 ratio). Within the training set, we used recursive feature elimination in a 5-fold cross validation setup to reduce the feature space (from 260,968 to 1805 oncRNA features) and identify a robust set of oncRNAs to use as our fingerprint. We then trained an XGBoost classifier with 500 trees on this set of 1805 oncRNAs to predict TOO on the whole training cohort. Applying the resulting model on the test data, we observed a strong performance with 90.9% accuracy. The performance metrics for each cancer are listed in **Fig S1I**, and the resulting confusion matrix reported in **Fig 1D** shows the fraction of samples of each cancer type that were correctly predicted. This confusion matrix is comparable to gene expression-based, genetic algorithm/k-nearest neighbors and convolutional neural network classifiers for TOO, including the higher number of mistakes in distinguishing rectal adenocarcinomas (READ) from colon adenocarcinoma (COAD), which were also found in other studies to be biological similar and often grouped together^11–13^. Interestingly, we also found that our model’s errors were enriched with misclassifications between different squamous cancers (*P* = 1.24 × 10^-13^, Fisher’s Exact Test), including bladder urothelial carcinoma (BLCA), cervical squamous cell carcinoma (CESC), esophageal carcinoma (ESCA), head and neck squamous cell carcinoma (HNSC), and lung squamous cell carcinoma (LUSC), consistent with previously reported unsupervised clusterings of different squamous tumors by various molecular platforms^14,15^. To emphasize the digital nature of oncRNA barcodes capable of distinguishing different cancer types, we plotted the binary expression patterns of oncRNAs selected by the XGBoost classifier for TOO classification (**Fig. S1J**). These results suggest that oncRNA expression patterns are informative of the underlying cancer biology, and thus our model can capture the heterogeneity of human cancers.

We also observed quantitative differences in the expression of oncRNAs beyond their binary presence-absence patterns, and thus asked whether including the relative oncRNA expression level could further improve our model’s TOO predictions (**Fig S1K, L**). To do this, we trained an XGBoost classifier using the counts per million (cpm)-based oncRNA expression profiles, using the same 80:20 ratio to split our samples into training and testing datasets. We found that the model trained on cpm data performed equally well with negligible differences and picked up important oncRNA features with similar patterns of expression as the binary model (**Fig S1M–Q**). The similarity in model performance of “digital” models trained on binarized oncRNA expression and “analog” models trained on normalized oncRNA expression data suggests that oncRNAs provide a digital barcode of cancer cell identity that is robust to the challenges in precise quantification of small RNA species.

Taken together, we have identified a large number of oncRNAs that are not only cancer-emergent but also reflective of cancer tissue-of-origin. We posited two likely routes for these orphan non-coding RNAs to emerge: (i) activation of cryptic promoters that lead to new transcriptional events and (ii) aberrant nucleolytic digestion of longer RNAs. We previously described T3p, a breast cancer-associated oncRNA derived from the *TERC* transcript, as an example of the latter pathway^1^. Mapping all of our newly identified oncRNAs to their genomic locations suggests that 58.9% of oncRNAs may originate from existing longer RNAs. In contrast, the 41.1% of oncRNAs that map to intergenic regions are more likely produced by cancer-specific transcriptional activation (**Fig S1R**). To explore this hypothesis further, we used roughly 386 ATAC-seq samples from TCGA to compare chromatin accessibility between samples as a function of oncRNA expression across tumors^16^. Approximately 10,000 intergenic oncRNA loci were captured at sufficient depth in the corresponding ATAC datasets. For a third of these loci, we observed a positive association between oncRNA expression in the small RNA data and chromatin accessibility in the ATAC-seq dataset, of which 1,989 oncRNA loci showed statistically significant associations at an FDR of 1% (**Fig 1E**). As expected, this association is entirely one-sided and we did not observe any oncRNAs in loci with closed chromatin. In **Fig 1F** and **Fig S1S**, we show the chromatin accessibility scores and relative expression of the top significant and expressed oncRNA loci as examples. This strong association between chromatin accessibility and oncRNA expression further supports our annotations and hypothesis that oncRNA biogenesis may arise from novel transcription events.

### oncRNA expression patterns are associated with cancer subtypes

In the previous section, we made two important observations: (i) oncRNAs show strong tissue-specific expression patterns and (ii) intergenic oncRNAs are associated with chromatin accessibility in cancer cells. Based on these findings, we hypothesized that oncRNA fingerprints may reflect the cellular state of cancer cells. To assess this possibility, we sought to identify oncRNAs whose presence or absence were informative of cancer subtypes. For this purpose, we used the Prediction Analysis of Microarray 50 (PAM50) breast cancer subtype classification (i.e., basal, HER2+, and luminal A and B) as well as the consensus molecular subtype (CMS) framework in colon cancer ^13,17^. Following the CMS classification system methodology, we combined the TCGA COAD and READ cohorts into a single colorectal cancer (CRC) cohort for all subsequent analyses ^13^. Of the 15,827 breast-cancer associated oncRNAs, 1,006 show significant subtype-specific patterns across the TCGA BRCA cohort **(Fig 2A)**.

**Figure 2.**
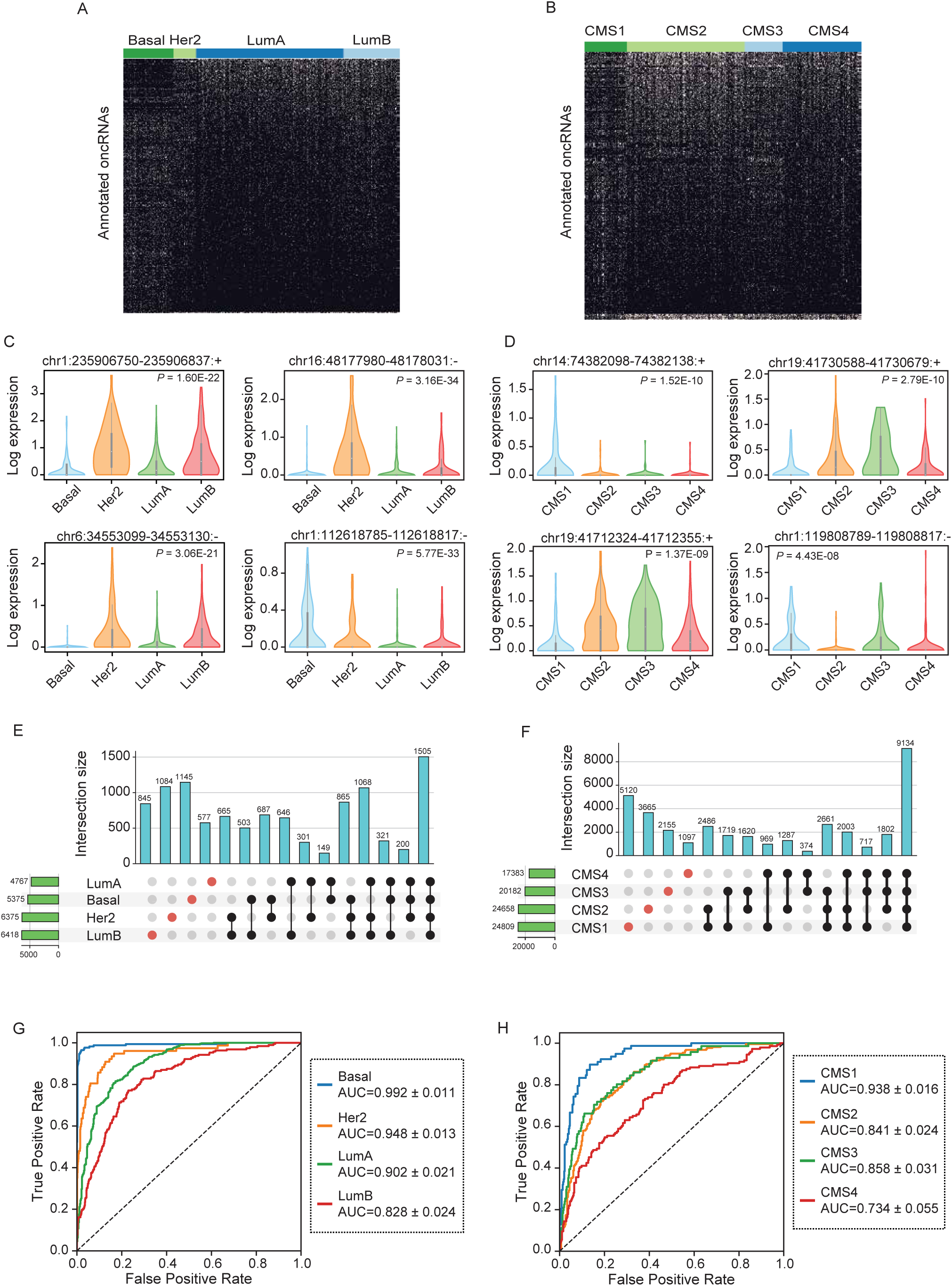
Annotation of subtype-associated oncRNAs across breast and colorectal cancer samples. **(A–B)** Binary heatmaps of oncRNAs associated with breast cancer subtypes (A) and colorectal cancer CMS labels (B). One-way ANOVA tests followed by FDR correction were used to identify oncRNAs with significant associations. **(C–D)** Exemplary subtype-associated oncRNA loci along with their expression patterns for breast cancer subtypes (C) or colon cancer CMS labels (D). The expression values are natural log transformed and *P* values were calculated using a one-way ANOVA test. **(E–F)** The number of oncRNAs that were detected in one or more breast cancer subtypes (E) or colorectal cancer CMS labels (F) shown as UpSet plots. **(G–H)** ROC curves for XGBoost multiclass classifiers that predict the breast cancer subtype or colon cancer CMS label based on oncRNA presence/absence fingerprints averaged across held-out validation sets in a 5-fold cross validation setup. 946 and 514 samples were tested in breast and colorectal cancer respectively and the resulting mean and standard deviation of AUCs were calculated for each subtype across the 5 folds.

For the TCGA CRC cohort, 1,198 of 57,632 CRC-associated oncRNAs demonstrate a significant association with CMS groups **(Fig 2B).** In **Fig 2C** and **2D**, we also included the normalized expression of several oncRNAs significantly associated with tumor subtypes after multiple testing correction (**Fig 2A, B**), highlighting the different quantitative patterns of expression across subtypes. Furthermore, we identified thousands of oncRNAs that were exclusively detected in samples of a given subtype for both breast and colorectal cancers, albeit insignificant when tested for subtype association across all samples **(Fig 2E, F).**

We then asked whether the cancer-associated oncRNAs could be leveraged to distinguish tumor subtypes using machine learning models. In a 5-fold cross-validation scheme, we used each training fold to train a multiclass XGBoost classifier. We then measured the performance of the model on the respective held-out fold. Breast cancer subtype classifications achieved AUCs between 0.83 and 0.99; similarly, colon cancer CMSs resulted in AUCs ranging between 0.73 and 0.94 (**Fig 2E–F)**. More detailed metrics of model performance for breast and colorectal cancers are reported in **Fig S2A** and **S2B**, respectively. Interestingly, we observed that the breast cancer model made a higher number of mistakes when distinguishing subgroups of luminal breast cancers, luminal A and luminal B, which are known to be more closely related and harder to distinguish^18^ (**Fig S2C**). We did not observe any notable patterns of confusion for CMS classification (**Fig 2SD).** We also show the binary patterns of all the oncRNA features selected by the XGBoost classifier within each training fold across all samples and the relative expression of the oncRNAs with the top 10 average feature importance score (**Fig S2E–H**). Our results indicate that the XGBoost model is able to learn and leverage a subset of informative oncRNAs from oncRNA fingerprints to accurately classify cancer subtypes for both breast and colorectal cancers. Together, these results further establish the utility of oncRNAs in not only distinguishing cancer tissue-of-origin, but also capturing their underlying cancer subtype identity.

### A systematic search for functional oncRNAs across multiple cancers

Given the regulatory potential of novel oncRNAs through oncRNA-RNA or oncRNA-protein interactions, we had previously investigated the possibility that oncRNAs may be adopted by cancer cells to engineer cancer-specific regulatory pathways^1^. Specifically, we uncovered one such oncRNA, T3p, and showed that it promotes breast cancer metastasis by dysregulating endogenous RISC complex activity. However, the extent to which other oncRNA species may play a functional role in cancer remains unexplored. The sheer number of oncRNA species emphasizes the need for systematic approaches to screen for functional representatives, in particular to identify oncRNAs that may drive tumorigenesis. To tackle this question, we developed a large-scale pooled *in vivo* screening framework to rapidly identify functional oncRNAs through gain- and loss-of-function studies. Our approach, schematized in **Fig 3A**, involves generating two libraries of lentiviral constructs: 1) a gain-of-function library encoding oncRNAs under the control of a U6 promoter to increase their expression; 2) a loss-of-function library of Tough Decoys (TuDs) to sequester oncRNAs, thereby inhibiting their endogenous functions^19^. To generate these libraries, we focused on four major cancers: breast, colon, lung, and prostate. We selected a human cell line with established xenograft models for each cancer: MDA-MB-231 for breast, SW480 for colon, A549 for lung, and C4-2B for prostate. We then used small RNA sequencing data from these cell lines to select expressed oncRNAs that were associated with each cell line’s respective tumor type in TCGA. For each cell line experiment, roughly 100 of the top expressed oncRNAs were selected for inclusion in the gain-of-function and loss-of-function (oncTuD) libraries. We also included non-targeting scramble sequences as endogenous controls. We transduced each of the four cell lines with their corresponding libraries and compared the representation of oncRNA/oncTuD species among cancer cell populations grown in mammary fat pads (MDA-MB-231) or subcutaneously (SW480, C4-2B, A549) *in vivo*, or grown *in vitro* for a similar number of doublings (**Fig S3A**). For each oncRNA/oncTuD instance, we compared their normalized counts between *in vivo* grown tumors and *in vitro* controls to identify those oncRNAs whose expression or TuD-mediated sequestration resulted in changes in the relative representation in the tumor context. We posited that changes in the baseline representation of cells harboring the cognate oncRNA or oncTuD lentiviral construct result from a selection pressure during tumorigenesis, which we can use as a criterion to identify functional oncRNAs.

**Figure 3.**
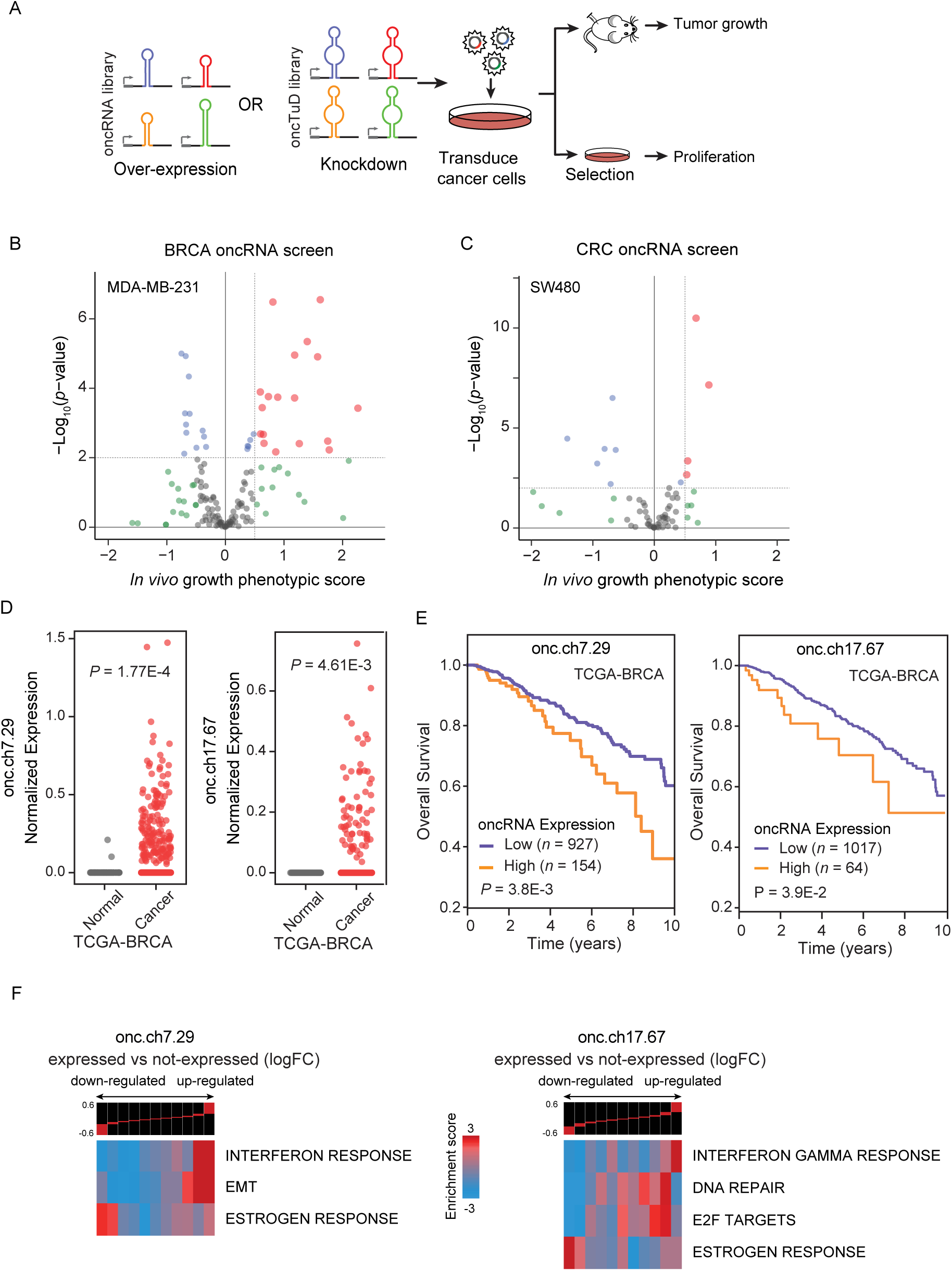
Systematic annotation of driver oncRNAs using a scalable *in vivo* genetic screening approach. **(A)** Workflow schematic of oncRNA cancer and oncRNA TuD functional screens. **(B-C)** Volcano plots of oncRNA functional screen results for breast cancer (MDA-MB-231) and colorectal cancer (SW480), respectively. *In vivo* growth phenotypic score refers to enriched representation of cancer cells transduced with cognate oncRNA upon tumor growth in the xenograft model. **(D)** Expression levels of two example oncRNAs with significant tumor growth phenotype from the functional screen in TCGA-BRCA tumor and tumor-adjacent normal tissues. *P* values were calculated using a one-tailed Mann-Whitney test. **(E)** Survival of TCGA-BRCA patients stratified by expression level of cognate driver oncRNA. *P* values were calculated using a log-rank test. **(F)** Informative iPage pathways associated with TCGA-BRCA cancer samples expressing cognate oncRNAs compared to TCGA-BRCA cancer samples with no detectable respective oncRNAs. Top panel shows gene expression differences in discrete expression bins. Genes that are up-regulated in oncRNA expressing cancer samples are in the right bins, whereas bins to the left contain genes with lower expression. The heatmap shows the corresponding pathway in relation to the expression bins. Red entries indicate enrichment of pathway genes in a given expression bin whereas blue entries indicate depletion. Enrichment and depletion are measured using log-transformed hypergeometric *P* values.

To identify oncogenic driver oncRNAs in our gain-of-function screens, we searched for those with increased expression in the tumors from the xenografted mice. We discovered several candidate functional oncRNAs in the breast and colon cancer screens; however, the lung and prostate cancer screens did not nominate any significant oncRNAs (**Fig 3B, C** and **Fig S3B**). Similarly, for the oncTuD screens, we selected oncRNAs whose antisense TuDs showed a reduced representation in the tumors. As shown in **Fig S3C–D**, a handful of oncRNAs showed a significant phenotypic effect within each cancer with the exception of breast cancer, which did not have oncRNAs with a significant phenotypic effect in the oncTuD screen. Our results indicate that between the gain and loss-of-function screens, roughly 5% of oncRNAs showed a significant tumor growth phenotype. This suggests that our earlier identification of T3p as a promoter of breast cancer metastasis was not a unique discovery and that cancer-emergent oncRNAs likely play unexplored roles in disease progression across human cancers. Together, these findings establish a systematic means of nominating likely functional oncRNA candidates impacting oncogenesis.

### Two oncRNAs that promote tumor growth and in vivo metastatic colonization of breast cancer cells

We next selected two exemplary breast cancer oncRNAs for a deeper analysis of their function. In **Fig 3D**, we compared the normalized expression levels of these two oncRNAs between TCGA-BRCA cancer and tumor-adjacent normal tissue samples and demonstrated the highly cancer-specific expression pattern of these oncRNAs (referred to by their respective genomic coordinates oncRNA.ch7.29 and oncRNA.ch17.67). Both oncRNA.ch7.29 and oncRNA.ch17.67 map to the 3’ UTRs of cancer-associated genes, *SCRN1* and *PSMD12* respectively. We also investigated the association of oncRNA expression with patient survival and found that these two oncRNAs were both significantly associated with poor clinical outcomes, further highlighting their potential functional role in breast cancer progression (**Fig 3E)**. However, we did not find any significant associations when we stratified oncRNA expression by cancer stage or receptor subtype for either oncRNA (**Fig S3E)**. To identify cellular processes and pathways that are associated with each of these two oncRNAs, we used the TCGA breast cancer dataset to compare the transcriptomic profiles between samples in which the oncRNA was detected versus those where it was not. We performed differential gene expression analysis and found significant changes in the gene expression landscape of tumors expressing each oncRNA (**Fig S3F**). Subsequent pathway analysis similarly revealed significant modulated pathways associated with the expression of each oncRNA, raising the possibility that they are acting downstream of these functional oncRNAs to drive cancer progression (**Fig 3F**, **S3G**)^20^. Of note, we observed a significant association between oncRNA.ch7.29 expression and up-regulation of genes in the EMT pathway, and significant associations between oncRNA.ch17.67 and up-regulation of genes in the DNA repair and E2F pathways.

We then performed *in vivo* tumor growth and metastasis assays to further validate the oncogenic role of these two oncRNAs. To test their effect, we first transduced MDA-MB-231 cells with oncRNA.ch7.29 or oncRNA.ch17.67 under the control of a U6 promoter for increased expression.

Overexpression of oncRNA.ch7.29 and oncRNA.ch17.67 both significantly increased the primary tumor growth rates of cells implanted in the mammary fat-pad of NOD *scid* gamma (NSG) mice by 2.6 and 1.7 folds, respectively, relative to scrambled controls **(Fig 4A)**. We then injected these transfected cells into the venous circulation of NSG mice and measured their lung metastatic colonization over time via bioluminescence imaging. Both oncRNA.ch7.29 and oncRNA.ch17.67 overexpressing cells had significantly increased capacity for lung colonization when compared to controls (**Fig 4B, S4A)**. We repeated these experiments in an independent breast cancer cell line, HCC1806 genetic background (HCC-LM2 ^21^), to ensure that our observations were not cell line dependent. We found that HCC-LM2 cells overexpressing oncRNA.ch7.29 or oncRNA.ch17.67 also exhibited significantly higher primary tumor rates and metastatic capacity (**Fig 4C–D, S4B)**.

**Figure 4.**
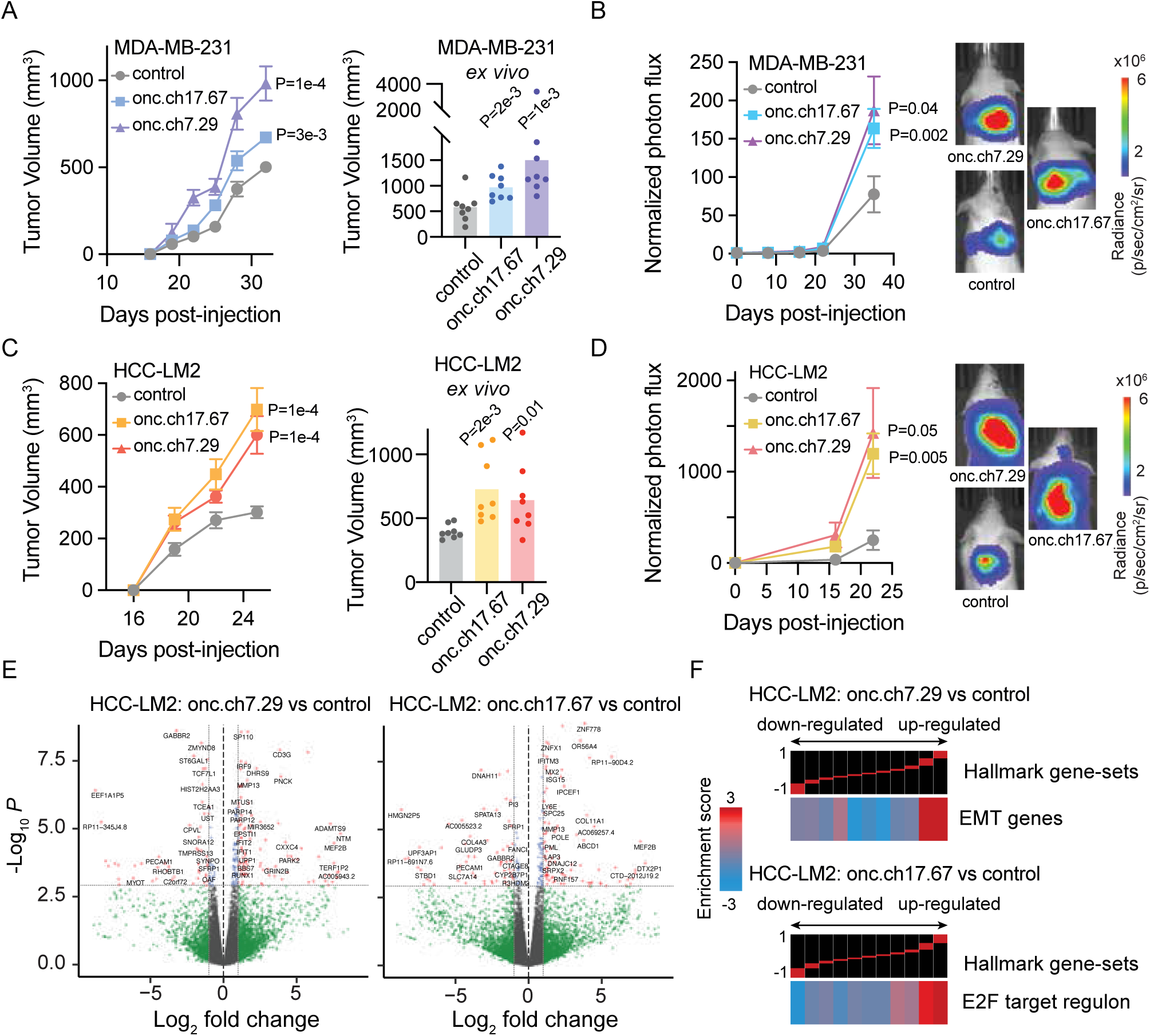
In vivo validation of functional oncRNAs in xenograft models of breast cancer. **(A)** Left: Growth of MDA-MB-231 tumors overexpressing oncRNA.ch7.29 or oncRNA.ch17.67 relative to controls in the mammary fat-pad of NSG mice. 2 tumors per mouse and n=4 mice for each cohort. *P* values were calculated using two-way ANOVA. Right: Ex vivo tumor measurements after tumor excision. *P* values were calculated using a one-tailed Mann-Whitney test. Tumors overexpressing oncRNA.ch7.29 were 2.6 fold larger than controls. Tumors overexpressing oncRNA.ch17.67 were 1.7 fold larger than controls. **(B)** Bioluminescence imaging plot of lung colonization by MDA-MB-231 cells overexpressing oncRNA.ch7.29 or oncRNA.ch17.67 compared to control. n = 5 per cohort. *P* values were calculated using two-way ANOVA. **(C)** Left: Growth of HCC-LM2 cells overexpressing oncRNA.ch7.29 or oncRNA.ch17.67 and HCC-LM2 controls in the mammary fat-pad of NSG mice mammary fat-pad assays. n=4 for each cohort. *P* values were calculated using two-way ANOVA. Right: Ex vivo tumor measurements after tumor excision. *P* values were calculated using a one-tailed Mann-Whitney test. Tumors overexpressing oncRNA.ch7.29 were 1.6 fold larger than controls. Tumors overexpressing oncRNA.ch17.67 were 1.8 fold larger than controls. **(D)** Bioluminescence imaging plot of lung colonization by HCC-LM2 cells overexpressing oncRNA.ch7.29 or oncRNA.ch17.67 compared to control. *n* = 5 per cohort. *P* values were calculated using two-way ANOVA. **(E)** Volcano plots of differentially expressed genes in HCC-LM2 cells overexpressing oncRNA.ch7.29 or oncRNA.ch17.67 compared to HCC-LM2 controls. The *P* value cut-off corresponds to a 10% FDR. **(F)** Representative pathways associated with HCC-LM2 overexpressing oncRNA.ch7.29 or oncRNA.ch17.67 compared to controls generated using iPAGE. Top panel shows gene expression differences in discrete expression bins. Genes that are up-regulated in oncRNA over-expressing cells are in the rightmost bins, whereas bins to the left contain genes with lower expression in oncRNA over-expressing cells. The heatmap shows the enrichment or depletion of the corresponding pathway in each expression bin. Red entries indicate enrichment of pathway genes in a given expression bin whereas blue entries indicate depletion.

We next asked if the function of these oncRNAs was mediated through the associated pathways we identified in TCGA-BRCA. To test this, we compared the transcriptomes of our cancer cells lines overexpressing oncRNA.ch7.29 or oncRNA.ch17.67 relative to controls in both genetic backgrounds (**Fig 4E, S4B)**. Pathway analysis of differential expression patterns revealed modulations in key oncogenic pathways that were also observed in our oncRNA association analysis in TCGA (Fig S4C– D), highlighting reproducible modulations of cellular pathways. Specifically, over-expressing oncRNA.ch7.29 resulted in an increase in the expression of epithelial-mesenchymal transition-related (EMT) genes, consistent with our observations in TCGA-BRCA tumors expressing oncRNA.ch7.29 (**Fig 4F, 3F).** Likewise, oncRNA.ch17.67 overexpressing cells demonstrated perturbation of the E2F pathway in a similar pattern as TCGA-BRCA tumors expressing oncRNA.ch17.67 (Fig 4F, 3F). While many significant oncRNA-associated pathways were shared among the HCC-LM2 and MDA-231 genetic backgrounds, we note that the E2F target regulon was not shown to be significantly associated with oncRNA.ch17.67 in MDA-231 cells (**Fig. S4D**). Together, our findings strongly support that a subset of oncRNAs drive oncogenesis, likely by perturbing specific gene pathways.

### Annotation of cell-free orphan non-coding RNAs across models of cancer

We have shown that oncRNA fingerprints represent a digital molecular barcode that effectively captures cancer type identity and are associated with modulations of cellular pathways that drive cancer progression. Importantly, oncRNA fingerprints have also shown the potential to be accessible from the extracellular space; we previously observed that a subset of breast cancer oncRNAs are secreted from breast cancer cells at detectable levels^1^. To investigate whether secreted oncRNA fingerprints are generalizable to other cancer types, we selected 25 established human cancer cell lines representing nine tissues of origin – blood, bone, breast, colon, kidney, lung, pancreas, prostate, and skin. After growing the cell lines *in vitro*, we collected conditioned media with exosome-depleted FBS in biological replicates, extracted RNA from the cell-free conditioned media, and performed smallRNA sequencing. It is known that many small RNAs, such as microRNAs, YRNAs, and tRNA fragments are secreted into the extracellular space ^22–25^. As shown in **Fig 5A–B** and **Fig S5A,** annotated small RNA profiles from biological replicates cluster together and, overall, cell lines from the same tissue of origin show similar patterns. We used this dataset of cell-free RNA content to identify oncRNAs that are expressed and secreted from each cell line. Overall, we observed cell-free small RNA reads mapping to thousands of oncRNA loci, making this biotype a significant contributor to the extracellular RNA space relative to other biotypes of smRNAs (**Fig 5C**). Roughly 0.5% of cell-free RNA reads were annotated as oncRNAs in our pipeline with about 30% of our pancancer list of oncRNAs detected in at least two cell lines (**Fig S5B**). Similar to our observation in tumor biopsies, we observed tumor type-specific oncRNAs among the cell-free oncRNAs (**Fig 5D**). Furthermore, UMAP visualization suggested an overall similarity between cell-free oncRNA fingerprints from cell lines of the same tumor type as their 2D UMAP projections clustered more closely together (**Fig 5E**). Similar clusterings of cell lines were also observed in the 2D PCA space of their oncRNA fingerprints (**Fig S5C**). To quantify this similarity, for each cell line, we compared the median correlation between its oncRNA profile with those from cell lines of the same tissue versus all other cell lines. Consistently, we observed a higher correlation between lines from the same tissue of origin than cell lines from different tissues of origin (**Fig S5D**).

**Figure 5.**
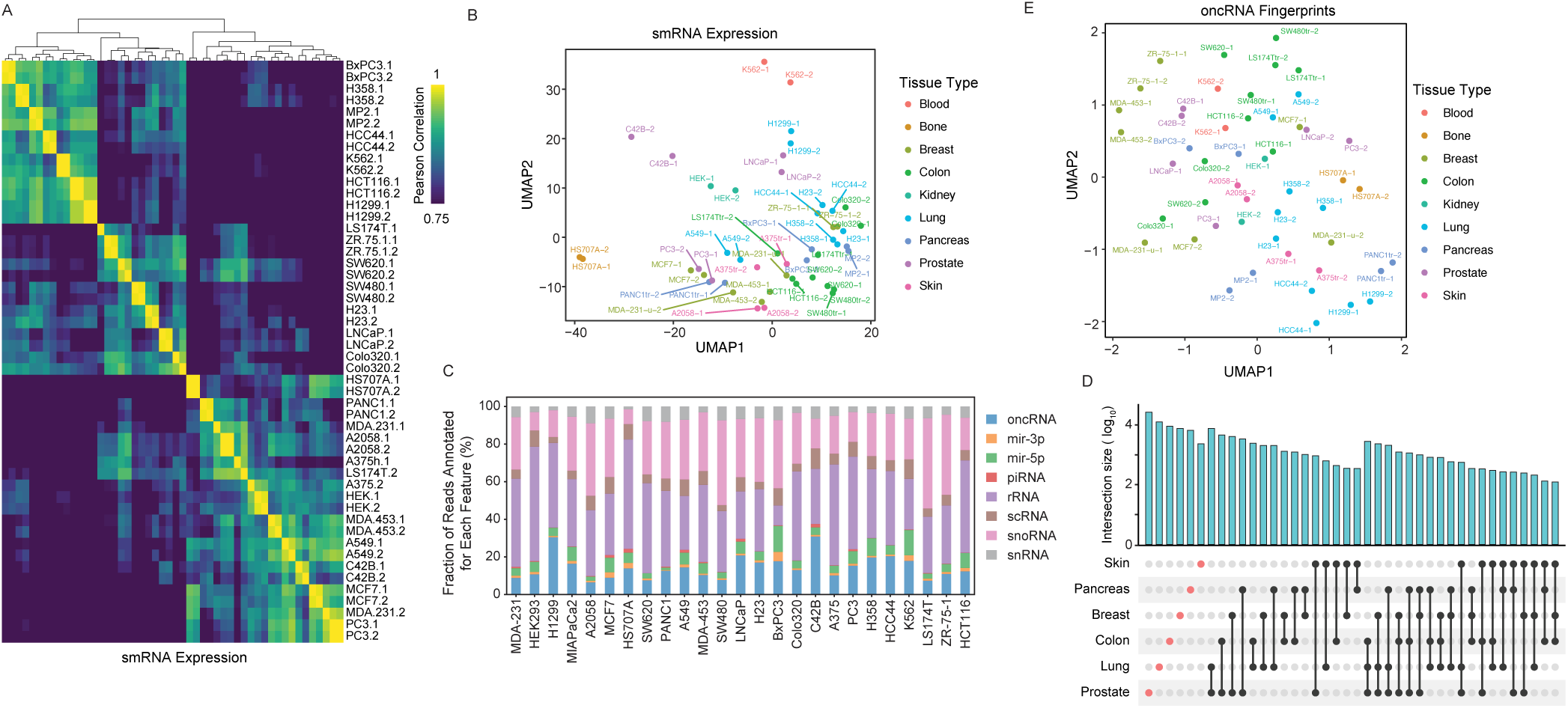
Analysis of cell-free RNA content across a large panel of cancer cell lines. **(A)** Pair-wise correlation heatmap for small RNA abundance in the cell-free RNA extracted from conditioned media. The counts for annotated small RNAs, such as miRNAs, tRNA fragments, snoRNAs, and *etc*, were used to generate this heatmap. **(B)** A 2D UMAP plot summarizing the abundance of small RNAs in the cell-free space across the cell line models we have profiled (in biological replicates). The points are colored based on the tissue-of-origin. **(C)** Contribution of each annotated family of small RNA species to their cell-free RNA content relative to annotated RNAs, omitting cell-free RNA with no known annotations. The values are normalized across cell lines and oncRNAs are shown in blue. **(D)** An UpSet plot of oncRNA counts detected in the cell-free RNA fraction of cell lines from each tissue-of-origin. Cell-free oncRNAs show tumor-specific patterns of expression. **(E)** 2D UMAP summary of oncRNA profiles across cell-free RNA profiles collected.

Taken together, our systematic analysis of cell-free RNA species secreted by cell line models of cancer demonstrates that oncRNAs contribute to the cell-free RNA content of cancer cells and that cell-free oncRNA expression profiles also reflect tumor type-specific patterns in these models.

### Circulating oncRNAs capture short-term and long-term clinical outcomes in breast cancer

Thus far, we have established that cell-free oncRNAs faithfully reflect cancer type identity. Since oncRNAs are cancer-emergent, their presence in circulation points to the presence of an underlying tumor that is actively releasing them. This notion is supported by our previous work showing that circulating T3p oncRNA can be used to detect breast cancer from serum in patients^1^. To assess the clinical utility of circulating oncRNAs as a cancer-specific biomarker, we performed a retrospective ancillary study on longitudinally collected samples from high-risk early breast cancer patients enrolled in the multicenter neoadjuvant I-SPY 2 TRIAL (NCT01042379)^26^. We extracted cell-free RNA from 1mL serum samples from 267 breast cancer patients treated in the I-SPY 2 TRIAL with standard neoadjuvant chemotherapy (NAC) alone or combined with MK-2206 (AKT inhibitor) or Pembrolizumab (PD-1 inhibitor) treatment. For each patient, we processed longitudinal serum samples collected at pretreatment (T0) and prior to surgery (T3) for small RNA sequencing. For 192 patients with T0 and T3 samples that passed our quality control filters, we measured total oncRNA burden, defined as the sum of all oncRNA species across all loci normalized by library size, for each time point. We then used the change in oncRNA burden before and after treatment (ΔoncRNA) as a measure of residual oncRNA burden. Detailed descriptions of our final patient cohort in our analysis are summarized in **Fig 6A** and **6SA**. In **Fig S6B**, we report the distribution of the resulting residual oncRNA burden classes across cancer subtypes, stages, and node status. Importantly, consistent with the response to treatment in the majority of patients, we observed a significant overall reduction in oncRNA burden after neoadjuvant chemotherapy (**Fig 6B, S6C**).

**Figure 6.**
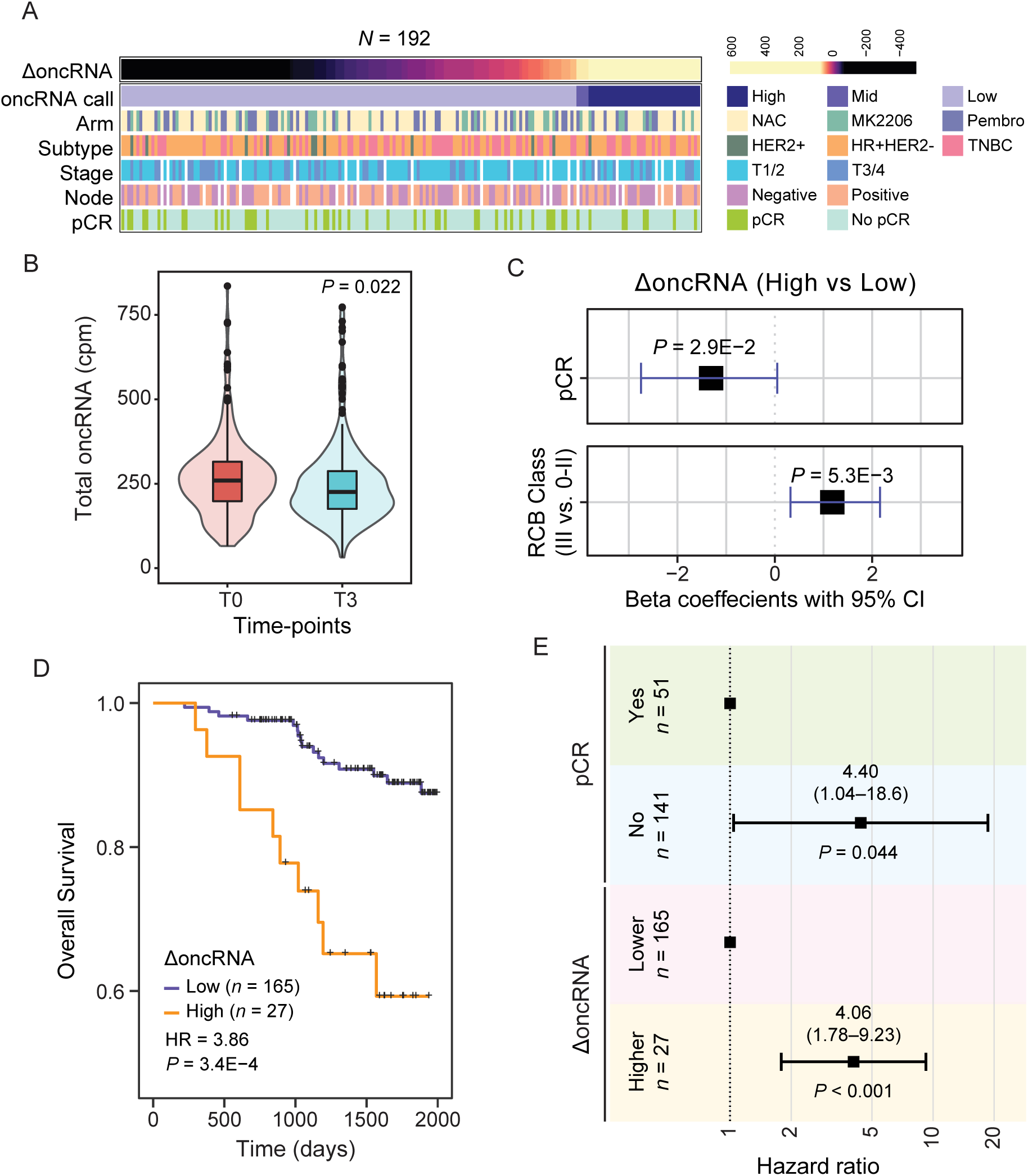
Changes in circulating oncRNA content over the course of neoadjuvant chemotherapy is informative of short-term and long-term clinical outcomes. **(A)** Overview of patient and tumor characteristics tabulated based on changes in oncRNA burden (ΔoncRNA). **(B)** Normalized oncRNA burden (counts per million) before (T0) and after (T3) neoadjuvant chemotherapy. *P* value was calculated using a one-tailed Wilcoxon test. **(C)** Forest plots for logistic regression models predicting pathologic complete response (pCR) or high residual cancer burden (RCB III) as a function of ΔoncRNA after neoadjuvant chemotherapy. One-tailed *P* values are also included. **(D)** Survival in patients grouped based on their oncRNA burden (ΔoncRNA). Reported are the hazard ratio and *P* value based on a log-rank test. **(E)** A forest plot for a multivariate Cox proportional hazard model including both ΔoncRNA and pCR as covariates.

Short-term clinical responses to NAC, i.e., pathologic complete response (pCR) and residual cancer burden (RCB) class, are strongly associated with favorable outcomes in the ISPY-2 trial. Thus, we first examined whether our ΔoncRNA calls were associated with these early clinical readouts. We used logistic regression to capture the association between high residual oncRNA burden after NAC with pCR and high RCB classification, respectively. As shown in **Fig 6C**, in both cases, we observed a significant association between residual oncRNA burden and short-term clinical responses.

With a median follow-up of 4.72 years in our study, we next sought to measure the extent to which residual oncRNA burden captures long-term clinical outcomes. For both overall survival and disease-free survival, we observed that high ΔoncRNA is significantly associated with poor survival outcomes (**Fig 6D, Fig S6E**). These associations were not highly sensitive to the choice of threshold for the high residual oncRNA burden call in patients (**Fig S6F**). Finally, we asked whether residual oncRNA burden provided additional information over pCR and RCB class regarding long-term survival. For this, we performed multivariable Cox regression analyses, and in both cases we observed that residual oncRNA burden remains significantly informative of survival even when controlling for pCR or RCB (**Fig 6E** and **Fig S6G**). Residual oncRNA burden also provided additional information when we controlled for tumor subtype and patient age (**Fig S6H)**, highlighting the limitations of subtyping in predicting treatment response and the added benefit of disease monitoring via oncRNA burden dynamics. These findings further highlight the tumor as the source of circulating oncRNAs in blood and establish these cell-free RNA species as clinically relevant liquid biopsy biomarkers that can be accessed from low volumes of blood.

## Discussion

In this study, we discovered and systematically annotated a previously unknown class of cancer-specific RNA species, oncRNAs, which have largely remained unexplored in the context of cancer biology. Our analysis not only reveals that these oncRNAs exhibit remarkable cancer type and subtype specificity, but also highlights the possible functional roles of oncRNAs for cancer progression.

Leveraging our *in vivo* screening platform, we revealed that a small subset of oncRNAs significantly impacts tumor growth phenotypes. Importantly, we view these numbers of significant hits to be a conservative estimate due to several factors. First, the lentiviral constructs described here are not guaranteed to up- or down-regulate their cognate oncRNAs: RNA Polymerase III-driven exogenous oncRNAs may be more unstable than endogenously expressed and processed oncRNAs, and TuDs may insufficiently inhibit their target oncRNAs, requiring fine-tuning of TuD design (i.e. optimizing thermodynamic properties) for adequate potency ^27^. Second, functions of oncRNAs are likely context-dependent, and the inclusion of other xenograft models will likely yield additional functional species. Finally, xenograft models only capture some aspects of tumor growth, lacking key characteristics such as adaptive immunity and native tumor microenvironment. Despite these limitations, our findings establish a systematic approach of combining *in vivo* screens and computational analysis to nominate new oncRNA drivers of oncogenesis. Namely, we consider oncRNAs that (i) display cancer-specific expression in both TCGA tumors and cancer cell line models, (ii) present a phenotypic effect in our functional screens, (iii) demonstrate significant association with poor clinical outcomes and (iv) cancer-relevant gene pathways as prime candidates for further functional or biogenesis investigations.

Although the molecular mechanism of action and biogenesis of oncRNA.ch7.29 and oncRNA.ch17.67 remains unknown, this study substantially expands our catalog of cancer-engineered oncogenic pathways and opens up exciting new avenues for exploring oncRNAs as novel therapeutic targets in cancer. Specifically, we found oncRNA.ch7.29 and oncRNA.ch17.67 to be significantly associated with modulations in EMT and E2F pathways, respectively. EMT is a crucial hallmark for cancer progression, particularly through loss of cell-adhesion, resistance to apoptosis, and acquired invasiveness ^28^. While non-coding RNAs like miRNAs have been shown to regulate cancer cell invasion and metastasis by targeting the mRNA of EMT-inducing transcription factors, our results suggest that cancer cells can also co-opt the complex EMT process via novel cancer-emergent RNA species ^29–31^.

The E2F target regulon collectively controls cell cycle progression, and are commonly activated in cancer cells to drive tumor proliferation ^32^. Consequently, there has been much attention for therapeutic interventions that affect E2F activity via targeting the CDK-RB-E2F axis throughCDK4/6 inhibitors for breast cancer ^33^. oncRNA.ch17.67’s upregulation of E2F genes may partially explain the increased tumor proliferation rate observed in our xenograft models and present as another potential therapeutic target to control E2F’s activity. Given E2F’s non-canonical role in apoptosis, metabolism, and angiogenesis, oncRNA.ch17.67 may also promote metastasis in a cell proliferation-independent manner ^32,34^. Because oncRNAs are largely absent in normal cells, targeting these cancer-associated pathways via oncRNAs may offer a specific therapeutic advantage by minimizing on-target toxicity and therefore reducing patient side effects.

Most importantly, our study shows that oncRNAs can be reliably detected in the circulating blood of cancer patients, making them valuable biomarkers for clinical applications. The current state-of-the-art liquid biopsy strategies for minimal residual disease detection in breast cancer rely on development of tumor-informed bespoke assays for detection of high variant allele frequency (VAF) mutations in the blood ^35,36^. Due to low DNA shedding from breast tumors, however, even with these bespoke assays DNA-based modalities are often not sensitive enough to reliably detect residual disease after clinical intervention^35^. Circulating oncRNAs allow us to overcome these limitations for liquid biopsy markers.

The much larger feature space of oncRNAs confers higher robustness against the zero-inflated nature of circulating biomarkers. Additionally, cancer cells actively secrete RNA; whereas DNA is passively shed as a result of cell death^37^. Thus, cell-free RNA biomarkers are often more abundant than their DNA counterparts, making oncRNAs highly sensitive biomarkers that can be detected even in low volumes of blood after treatment. Other cell-free RNAs, including microRNAs, repeat element derived RNAs, and transfer RNA-derived small RNAs, have also been of recent research interest for their potential as circulating biomarkers of cancer^38–41^. While prior studies have shown cfRNA profiles to be promising for applications in cancer detection, cfRNA signatures have primarily been discovered directly from human plasma samples and are unlikely to be directly representative of the underlying tumor biology or state. These signatures also predominantly rely on RNAs of known annotations that can originate from any cell and may not be directly secreted by cancer cells. Furthermore, investigations of cfRNAs as clinical biomarkers have largely been restricted to applications in cancer detection with limited success.

In our retrospective study, we investigated the utility of circulating oncRNAs for minimum residual disease detection and predicting clinical outcome in a neoadjuvant chemotherapy setting. We combined all oncRNA species to define an oncRNA burden score and found the dynamic changes in the oncRNA burden score in response to neoadjuvant chemotherapy to be strongly associated with both short-term clinical responses and long-term survival outcomes. These results establish oncRNAs as biomarkers for minimally-invasive and real-time monitoring of underlying cancers, which can significantly help guide cancer management. We anticipate that future liquid biopsy studies with substantially larger cohort sizes as well as larger collected blood volumes and deeper sequencing of the cell-free RNA content will enable us to delve deeper into the wealth of information offered by oncRNAs and potentially reveal new cancer-subtype signatures, cancer subtype switching occurrences, or relationships to treatment response.

In conclusion, our study has unveiled a previously unannotated class of RNA species, oncRNAs, which hold immense potential for both disease monitoring and therapeutic applications in cancer. As we continue to investigate the various roles and information carried out by individual oncRNAs, we anticipate that these RNA species will prove to be invaluable tools in the ongoing battle against cancer.

## Acknowledgements

The I-SPY2 trial is supported by the study sponsors, Quantum Leap Healthcare Collaborative (2013 to present) and a grant from the National Cancer Institute (P01CA210961). Sequencing was performed at the UCSF CAT, supported by UCSF PBBR, RRP IMIA, and NIH 1S10OD028511-01 grants. Friends for breast cancer, AACR, Emerson Collective, Mark Foundation. HG is an Era of Hope Scholar (W81XWH-2210121) and supported by grants from NCI (R01CA240984 and R01CA244634). This work was also supported by Earlier, the Emerson Collective, and the Mark Foundation. We thank Amy L. Delson and Carol Simmons, our patient advocates, for their valuable feedback throughout this study.

## Disclosure of Potential Competing Interest

H.G. is a co-founder and shareholder of Exai Bio. J.W., L.F., and T.C. are employees and shareholders of Exai Bio. L.J.E. reports funding from Merck & Co.; participation on an advisory board for Blue Cross Blue Shield; and personal fees from UpToDate. L.J.v.V. is a founding advisor and shareholder of Exai BIo; part-time employee and owns stock in Agendia. All other authors declare no competing interests.

