## Supplemental Figs and Methods for "Systematic annotation of orphan RNAs reveals blood-accessible molecular barcodes of cancer identity and cancer-emergent oncogenic drivers"

### Supplementary Figures

A

| Project | Biofluid | Number of Samples |
|---|---|---|
| GSE112343 | Bile | 12 |
| GSE113994 | Plasma | 212 |
| | Serum | 94 |
| GSE121978 | Urine | 78 |
| GSE123336 | Saliva | 87 |
| | Serum | 131 |
| GSE128348 | Urine | 205 |
| GSE128359 | Plasma | 123 |
| | Stool | 33 |
| | Urine | 47 |
| GSE140069 | Blood | 105 |
| GSE46579 | Blood | 69 |
| GSE53439 | Blood | 2 |
| | Serum | 22 |
| phs001258 | Plasma | 180 |
| | Saliva | 36 |
| | Urine | 203 |

B

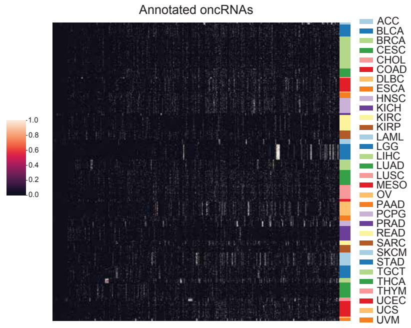


C

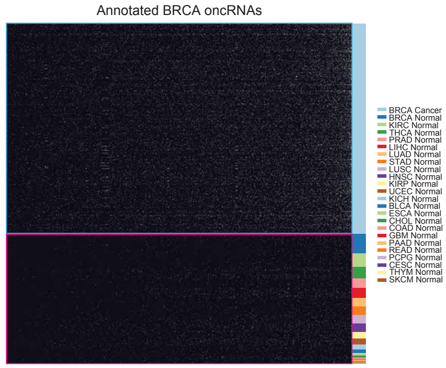


D

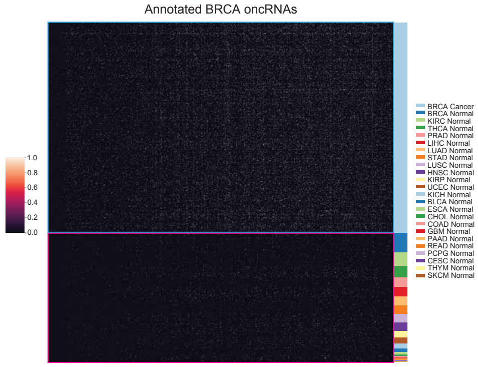


E

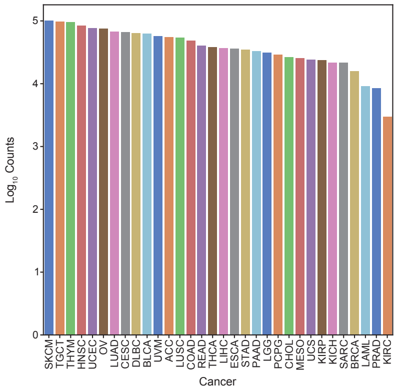


F

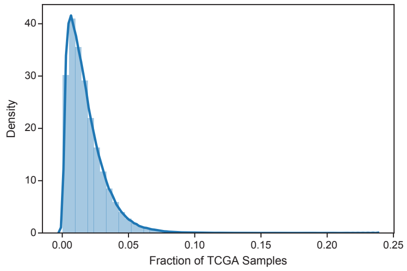


G

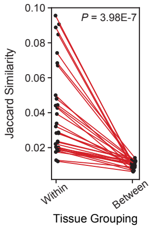


H

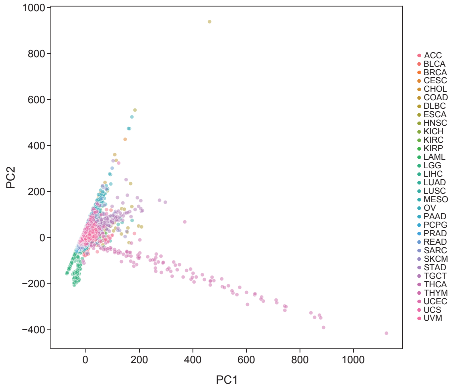

I

| Cancer | Precision | Recall | f1-Score |
|---|---|---|---|
| ACC | 0.94 | 0.94 | 0.94 |
| BLCA | 0.81 | 0.79 | 0.80 |
| BRCA | 0.90 | 0.98 | 0.94 |
| CESC | 0.86 | 0.69 | 0.77 |
| CHOL | 1.00 | 0.86 | 0.92 |
| COAD | 0.80 | 0.89 | 0.85 |
| DLBC | 0.88 | 0.78 | 0.82 |
| ESCA | 0.96 | 0.65 | 0.77 |
| HNSC | 0.84 | 0.88 | 0.86 |
| KICH | 1.00 | 0.92 | 0.96 |
| KIRC | 0.91 | 0.97 | 0.94 |
| KIRP | 0.95 | 0.91 | 0.93 |
| LAML | 1.00 | 0.97 | 0.99 |
| LGG | 1.00 | 1.00 | 1.00 |
| LIHC | 0.99 | 0.93 | 0.96 |
| LUAD | 0.89 | 0.91 | 0.90 |
| LUSC | 0.72 | 0.81 | 0.76 |
| MESO | 0.94 | 0.88 | 0.91 |
| OV | 0.99 | 0.99 | 0.99 |
| PAAD | 0.94 | 0.83 | 0.88 |
| PCPG | 1.00 | 0.97 | 0.99 |
| PRAD | 1.00 | 0.97 | 0.98 |
| READ | 0.58 | 0.44 | 0.50 |
| SARC | 0.87 | 0.89 | 0.88 |
| SKCM | 0.98 | 0.99 | 0.98 |
| STAD | 0.89 | 0.89 | 0.89 |
| TGCT | 1.00 | 0.97 | 0.98 |
| THCA | 0.97 | 0.94 | 0.96 |
| THYM | 1.00 | 1.00 | 1.00 |
| UCEC | 0.89 | 0.93 | 0.91 |
| UCS | 1.00 | 0.82 | 0.90 |
| UVM | 1.00 | 0.94 | 0.97 |
| **Accuracy** | | | 0.91 |
| **Macro-average** | 0.92 | 0.89 | 0.90 |
| **Weighted-average** | 0.91 | 0.91 | 0.91 |

J

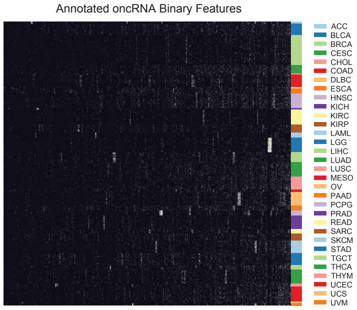


K

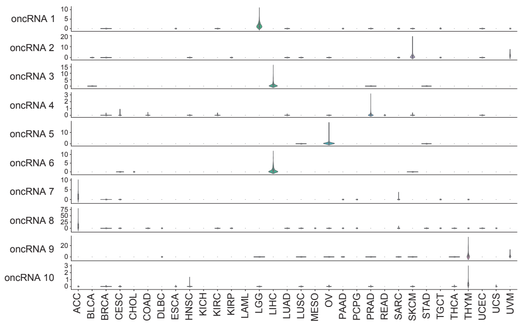


L

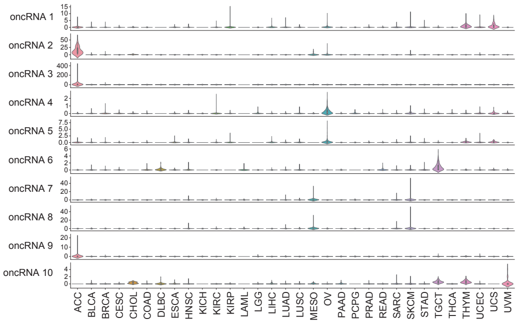


M

| Cancer | Precision | Recall | f1-Score |
|---|---|---|---|
| ACC | 0.94 | 1.00 | 0.97 |
| BLCA | 0.88 | 0.77 | 0.82 |
| BRCA | 0.90 | 0.98 | 0.94 |
| CESC | 0.82 | 0.79 | 0.80 |
| CHOL | 0.86 | 0.86 | 0.86 |
| COAD | 0.81 | 0.87 | 0.84 |
| DLBC | 0.75 | 0.67 | 0.71 |
| ESCA | 0.88 | 0.59 | 0.71 |
| HNSC | 0.79 | 0.89 | 0.84 |
| KICH | 1.00 | 0.92 | 0.96 |
| KIRC | 0.93 | 0.97 | 0.95 |
| KIRP | 0.97 | 0.97 | 0.97 |
| LAML | 0.97 | 0.97 | 0.97 |
| LGG | 1.00 | 1.00 | 1.00 |
| LIHC | 0.99 | 0.93 | 0.96 |
| LUAD | 0.84 | 0.88 | 0.86 |
| LUSC | 0.76 | 0.82 | 0.79 |
| MESO | 1.00 | 0.82 | 0.90 |
| OV | 0.99 | 0.99 | 0.99 |
| PAAD | 1.00 | 0.83 | 0.91 |
| PCPG | 1.00 | 0.97 | 0.99 |
| PRAD | 1.00 | 0.98 | 0.99 |
| READ | 0.58 | 0.47 | 0.52 |
| SARC | 0.88 | 0.96 | 0.92 |
| SKCM | 0.98 | 0.99 | 0.98 |
| STAD | 0.85 | 0.81 | 0.83 |
| TGCT | 1.00 | 0.97 | 0.98 |
| THCA | 1.00 | 0.93 | 0.96 |
| THYM | 1.00 | 1.00 | 1.00 |
| UCEC | 0.89 | 0.90 | 0.89 |
| UCS | 1.00 | 0.73 | 0.84 |
| UVM | 1.00 | 0.94 | 0.97 |
| **Accuracy** | | | 0.91 |
| **Macro-average** | 0.91 | 0.88 | 0.89 |
| **Weighted-average** | 0.91 | 0.91 | 0.91 |

N

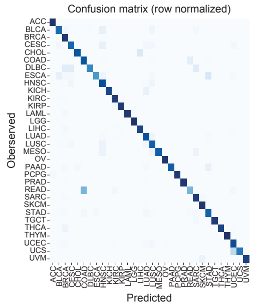


O

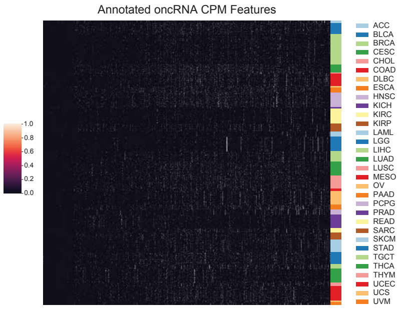

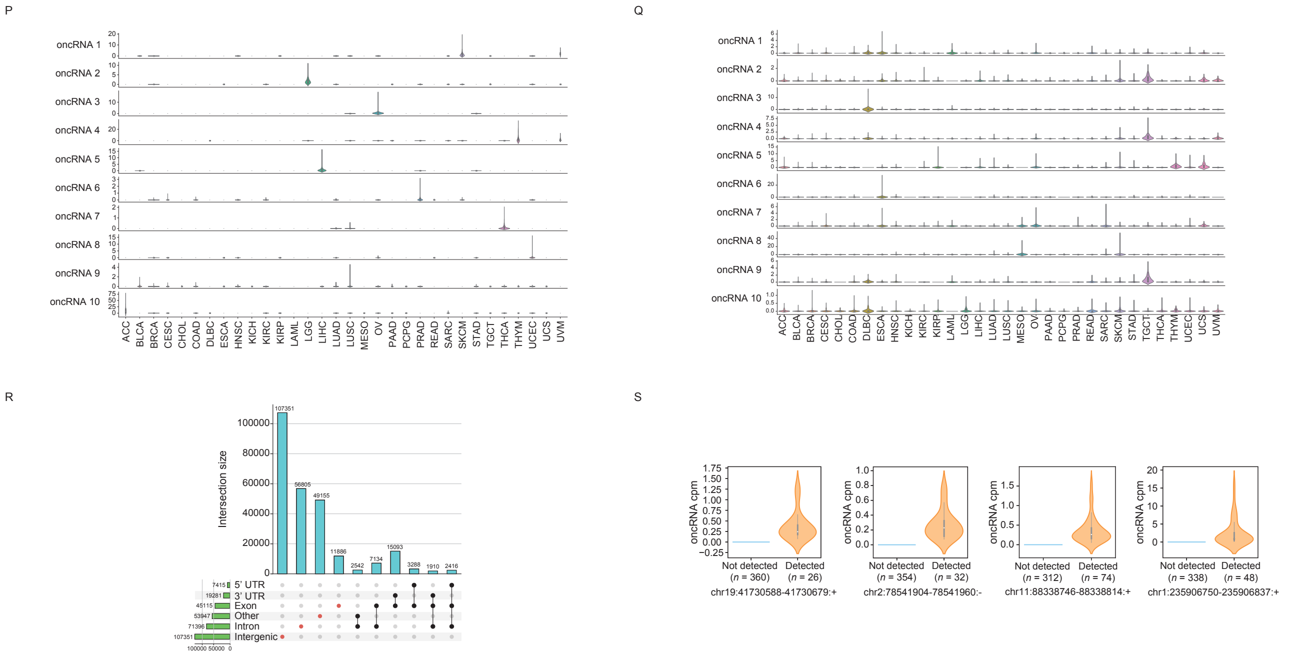


**Supplementary Figure 1. Discovery and profiling of oncRNAs in cancer tissues.**

**(A)** Table of publicly available datasets from the exRNA Atlas used to filter RNAs. **(B)** A heat map of normalized expression of oncRNAs across TCGA samples. Each row represents a sample and each column represents an oncRNA. **(C-D)** Binary and normalized expression heatmap of oncRNAs annotated in the TCGA-BRCA cohort, respectively. A total of 16,474 breast cancer oncRNAs were annotated and plotted here. **(E)** $Log_{10}$ number of oncRNAs annotated in each cancer type. **(F)** Density plot of the fraction of TCGA samples for which each of the 260,968 onRNAs was observed. **(G)** Median Jaccard similarity of oncRNA profiles between cancer samples from the same cancer tissue group versus different cancer tissue groups. *P* value was calculated using a one-tailed Wilcoxon test. **(H)** PCA plot of oncRNA profiles of all TCGA cancer samples. Points are colored by the cancer types. **(I)** Performance metrics of the tissue-of-origin (TOO) XGBclassifer trained on binary oncRNA profiles and evaluated on the held-out dataset. **(J)** Binary heatmap of the oncRNAs used as binarized features for the TOO XGBclassifier model. **(K)** Expression levels of top 10 important and **(L)** prevalent oncRNAs in the TOO XGBclassifier model trained on binary oncRNA profiles. Ranking of oncRNA feature importance is based on average information gain as determined by the model during training. **(M)** Performance metrics of the final XGBclassifer trained on normalized oncRNA expression profiles (counts-per-million) and evaluated on the held-out dataset. **(N)** The confusion matrix for tissue-of-origin classification based on normalized expression of oncRNAs in each sample. The matrix was row-normalized. **(O)** Heatmap of the normalized expression of oncRNAs used as features for the TOO XGBclassifier model in (L). **(P)** Expression levels of top 10 important and **(Q)** prevalent oncRNAs in the TOO XGBclassifier model trained on normalized oncRNA expression profiles. **(R)** Upset plot depicting the overlaps of oncRNAs with established genomic features. The other category refers to overlaps of oncRNAs to the opposite strand of the genomic features. oncRNAs with no overlaps with the genomic features were placed in the intergenic category. **(S)** Normalized expression levels of four exemplary oncRNAs. Expression level of cognate oncRNA was used to split samples into detected and not detected groups for the chromatin accessibility analysis (Fig 1F). Values are shown as violin plots and boxplots. The boxplots show the distribution quartiles, and the whiskers show the quartiles ± IQR (interquartile range). Also reported are the number of samples in which the oncRNAs were detected.

A

| BRCA Subtype | Precision | Recall | f1-Score |
|---|---|---|---|
| Basal | 0.93 (0.072) | 0.97 (0.013) | 0.94 (0.044) |
| Her2 | 0.79 (0.144) | 0.62 (0.148) | 0.69 (0.114) |
| LumA | 0.80 (0.016) | 0.89 (0.029) | 0.84 (0.010) |
| LumB | 0.58 (0.054) | 0.42 (0.061) | 0.48 (0.037) |
| Accuracy | | | 0.78 (0.017) |
| Macro-average | 0.77 (0.037) | 0.72 (0.039) | 0.74 (0.035) |
| Weighted-average | 0.77 (0.019) | 0.78 (0.017) | 0.77 (0.018) |

B

| CRC Subtype | Precision | Recall | f1-Score |
|---|---|---|---|
| CMS1 | 0.69 (0.083) | 0.65 (0.149) | 0.66 (0.083) |
| CSM2 | 0.68 (0.033) | 0.79 (0.032) | 0.73 (0.030) |
| CMS3 | 0.56 (0.124) | 0.42 (0.082) | 0.48 (0.091) |
| CMS4 | 0.53 (0.092) | 0.47 (0.088) | 0.49 (0.083) |
| Accuracy | | | 0.63 (0.037) |
| Macro-average | 0.61 (0.044) | 0.58 (0.046) | 0.59 (0.044) |
| Weighted-average | 0.62 (0.039) | 0.63 (0.037) | 0.62 (0.039) |

C

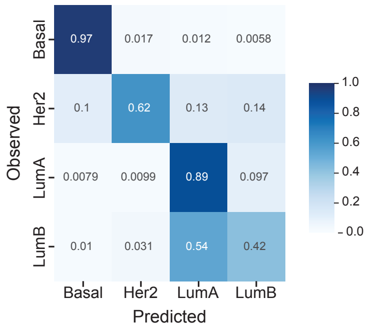


D

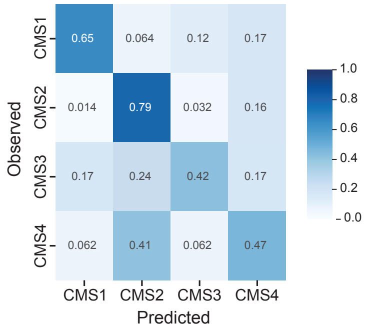


E

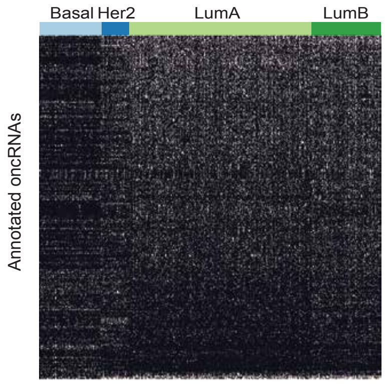


F

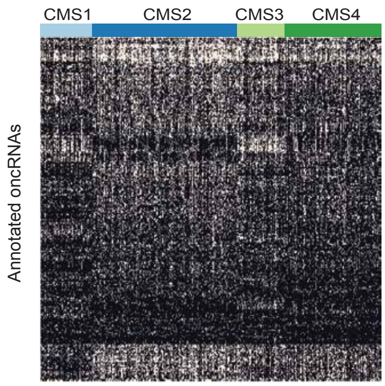


G

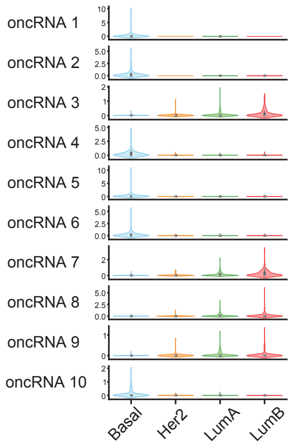


H

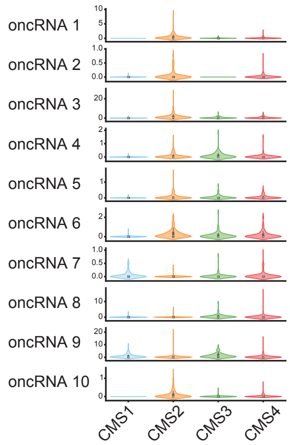

**Supplementary Figure 2. Analysis of subtype specific oncRNAs in breast and colorectal cancers.**
**(A)** Performance metrics of the breast cancer subtype XGBclassifer averaged (standard deviation) across 5 folds. **(B)** Performance metrics of the colorectal cancer subtype XGBclassifer averaged (standard deviation) across 5 folds. **(C)** The confusion matrix for breast cancer subtype classification averaged across 5 folds for the XGBclassifier. The matrix was row-normalized. **(D)** The confusion matrix for colorectal cancer subtype classification averaged across 5 folds for the XGBclassifier. The matrix was row-normalized. **(E–F)** Binary heatmap of oncRNAs used as features by the XGBclassifier for breast cancer (E) and colorectal cancer (F). **(G–H)** Expression levels of top 10 important oncRNAs in the XGBclassifier models trained on binary oncRNA expression profiles to predict breast cancer subtype (G) and colorectal cancer subtype (H).

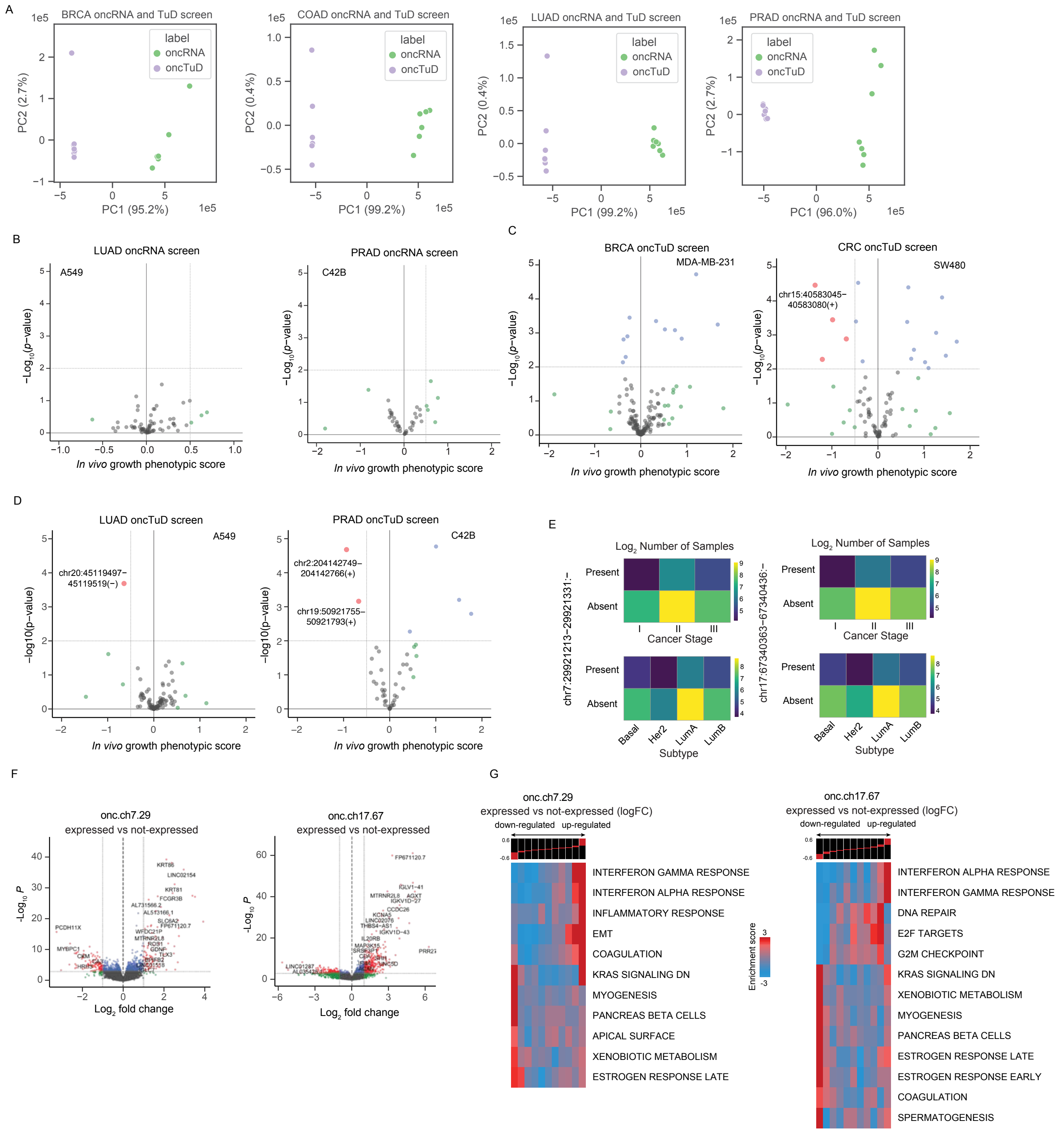
A
BRCA oncRNA and TuD screen
COAD oncRNA and TuD screen
LUAD oncRNA and TuD screen
PRAD oncRNA and TuD screen
label
oncRNA
oncTuD
PC1 (95.2%)
PC2 (2.7%)
PC1 (99.2%)
PC2 (0.4%)
PC1 (96.0%)
B
LUAD oncRNA screen
A549
PRAD oncRNA screen
C42B
In vivo growth phenotypic score
C
BRCA oncTuD screen
MDA-MB-231
CRC oncTuD screen
SW480
chr15:40583045–40583080(+)
D
LUAD oncTuD screen
A549
chr20:45119497–45119519(−)
PRAD oncTuD screen
C42B
chr2:204142749–204142766(+)
chr19:50921755–50921793(+)
E
Log2 Number of Samples
Present
Absent
Cancer Stage
Subtype
Basal
Her2
LumA
LumB
chr7:29921213–29921331:−
chr17:67340363–67340436:−
F
onc.ch7.29 expressed vs not-expressed
onc.ch17.67 expressed vs not-expressed
Log2 fold change
G
onc.ch7.29 expressed vs not-expressed (logFC)
onc.ch17.67 expressed vs not-expressed (logFC)
down-regulated
up-regulated
Enrichment score
INTERFERON GAMMA RESPONSE
INTERFERON ALPHA RESPONSE
INFLAMMATORY RESPONSE
EMT
COAGULATION
KRAS SIGNALING DN
MYOGENESIS
PANCREAS BETA CELLS
APICAL SURFACE
XENOBIOTIC METABOLISM
ESTROGEN RESPONSE LATE
DNA REPAIR
E2F TARGETS
G2M CHECKPOINT
ESTROGEN RESPONSE EARLY
SPERMATOGENESIS

**Supplementary Figure 3. *In vivo* screen to identify oncRNAs with functional roles during cancer progression.**

**(A)** PCA plot of oncRNA and oncRNA Tough Decoy (oncTuD) expression in breast (BRCA; MDA-MB-231), colorectal (CRC; SW480), lung (LUAD; A549), and prostate (PRAD; C4-2B) cancer cell lines transduced with cognate oncRNA (green) or oncTuD (purple) libraries. Each cancer gain-of-function and loss-of-function screen was done in replicates. **(B)** Volcano plots of onRNA functional screen results for lung cancer (A549) and prostate cancer (C42B), respectively. *In vivo* growth phenotypic score refers to enriched representation of cancer cells transduced with cognate oncRNA upon tumor growth in the xenograft model. **(C–D)** Volcano plots of onRNA TuD functional screen results for breast cancer (MDA-MB-231) and colorectal cancer (SW480) (C) and lung cancer (A549) and prostate cancer (C42B) (D). **(E)** $Log_2$ count matrices of TCGA breast cancer samples stratified by cancer stage (top) or subtype (bottom) for which two driver oncRNAs with significant tumor growth phenotype were present or absent. **(F)** Volcano plots of differentially expressed genes in TCGA-BRCA tumors expressing the specified oncRNA compared with tumors in which cognate oncRNA was undetected. **(G)** Full list of informative iPage pathways associated with TCGA-BRCA tumors expressing cognate oncRNAs compared to TCGA-BRCA tumors in which respective oncRNAs were not detected.

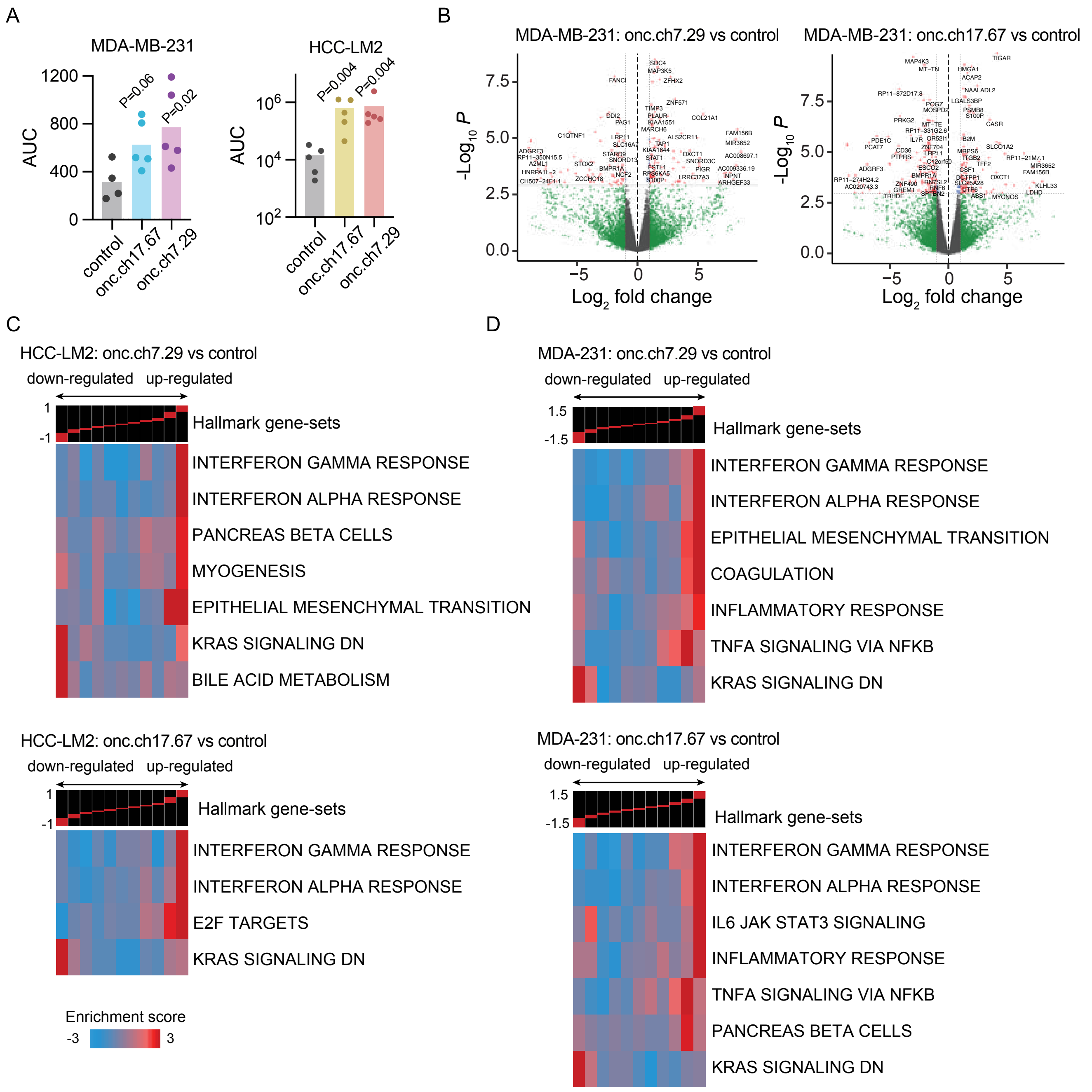


**Supplementary Figure 4. Validation of function oncRNAs *In vivo* models of cancer progression.**
**(A)** Area under the curve (AUC) of the bioluminescence plots from the lung colonization assays with MDA-MB231 cell lines (left) and HCC-LM2 cell lines (right), corresponding with Fig 4B and 4D, respectively. *P* values were calculated using a one-tailed Mann-Whitney test. **(B)** Volcano plots of differentially expressed genes in MDA-MB231 cells overexpressing oncRNA.ch7.29 or oncRNA.ch17.67 compared to MDA-MB231 controls. **(C–D)** Informative iPage pathways associated with HCC-LM2 cells overexpressing oncRNA.ch7.29 or oncRNA.ch17.67 compared to controls (C) and MDA-231 cells overexpressing oncRNA.ch7.29 or oncRNA.ch17.67 compared to controls (D).

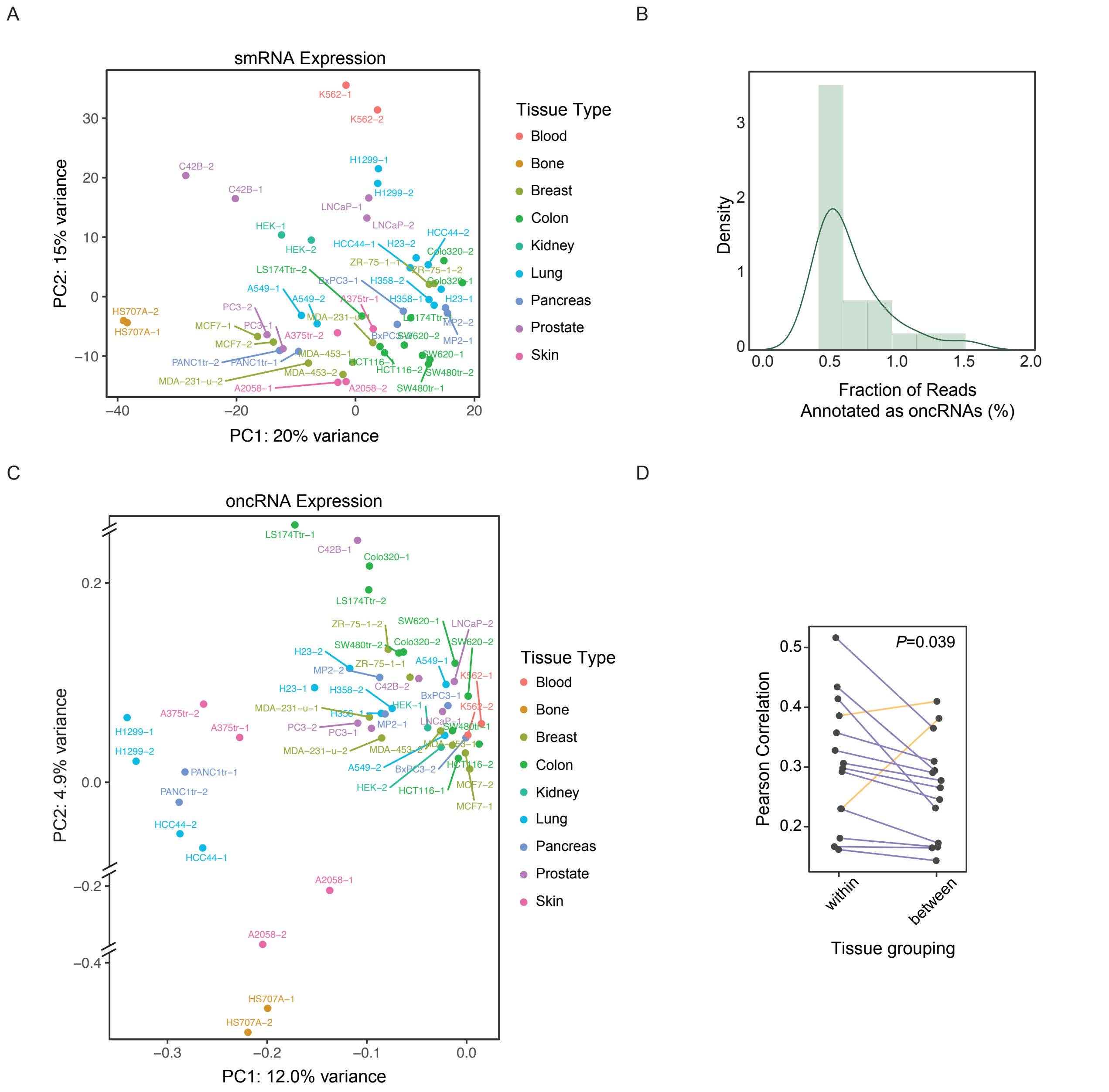


**Supplementary Figure 5. oncRNAs reflect cancer cell line identity in extracellular space.**

**(A)** PCA plot of the cell-free smRNA expression profiles of 25 cancer cell lines. Points are colored by the tumor type of the cell lines. **(B)** Density plot of the fraction of reads annotated as oncRNAs across all cancer cell lines in (A). **(C)** PCA plot of the cell-free oncRNA expression profiles in the cancer cell lines. Points are colored by the cell lines' corresponding tumor types. **(D)** Median Pearson correlation of oncRNA profiles between cell lines of the same tumor type (within) and cell lines of the same tumor type versus all other cell lines (between). Each connected pair of points consists of one reference tumor type. Tumor types with higher between-cancer tissue group correlations are colored orange, while tumor types with higher within group correlations are colored purple. Also reported is the *P* value calculated using a two-tailed paired Student's *t*-test.

A

| | | Missing | Overall |
|---|---|---|---|
| ***N*** | | | 192 |
| **Age, mean (SD)** | | 0 | 47.9 (10.9) |
| **T Stage, *n* (%)** | T1/T2 | 0 | 105 (54.7) |
| | T3/T4 | | 67 (34.9) |
| | Undefined | | 20 (10.4) |
| **Node Status, *n* (%)** | Node+ | 0 | 89 (46.4) |
| | Node- | | 78 (40.6) |
| | Undefined | | 25 (13.0) |
| **Subtype, *n* (%)** | HER2+ | 0 | 12 (6.2) |
| | HR+HER2- | | 109 (56.8) |
| | TNBC | | 71 (37.0) |
| **pCR, *n* (%)** | 0 | 0 | 141 (73.4) |
| | 1 | | 51 (26.6) |
| **RCB, *n* (%)** | 0 | 0 | 53 (27.6) |
| | I | | 27 (14.1) |
| | II | | 79 (41.1) |
| | III | | 29 (15.1) |
| | Undefined | | 4 (2.1) |
| **Arm, *n* (%)** | Pac | 0 | 105 (54.7) |
| | Pac + MK | | 25 (13.0) |
| | Pac + MK + Tras | | 10 (5.2) |
| | Pac + Pemb | | 50 (26.0) |
| | Pac + Tras | | 2 (1.0) |
| **oncRNA, *n* (%)** | High | 0 | 37 (19.3) |
| | Mid | | 4 (2.1) |
| | Low | | 151(78.6) |

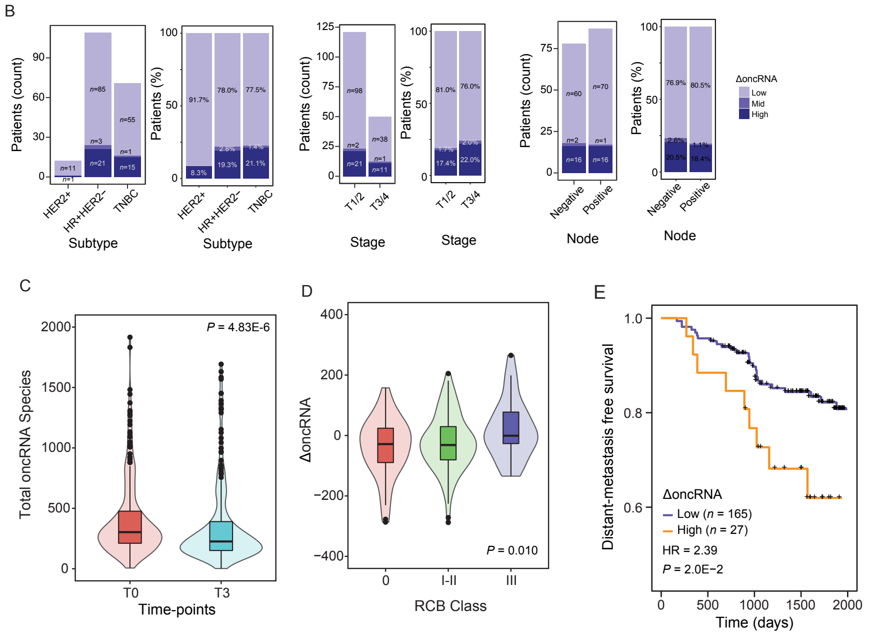


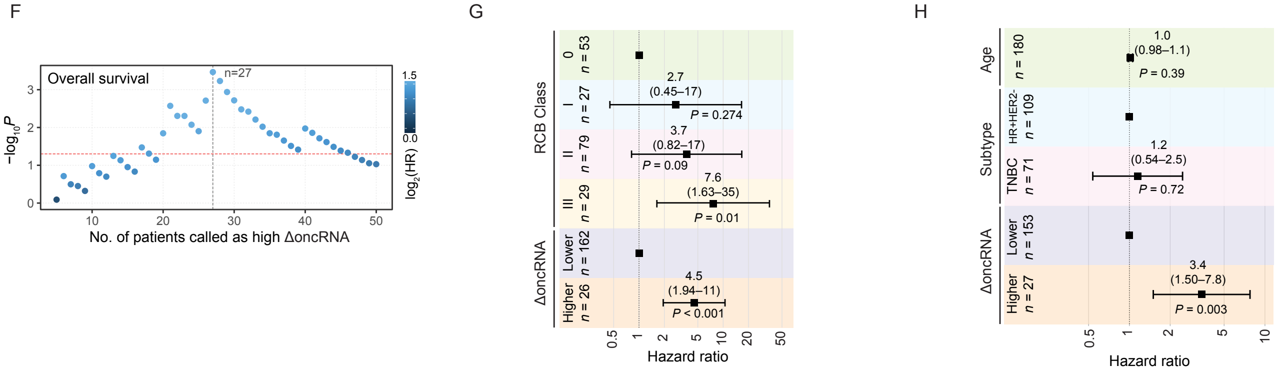


**Supplementary Figure 6. Analysis of residual oncRNA burden in the ISPY-2 trial cohort.**

**(A)** Summary statistics of the ISPY-2 trial patient cohort (*n* = 192). Only patients with samples that passed our quality control filters for both time point 0 (prior to neoadjuvant chemotherapy) and time point 3 (prior to surgery) are included in this table. **(B)** Distributions of residual oncRNA burden (ΔoncRNA) levels among ISPY-2 patients, grouped by breast cancer subtype, tumor T classification, and node status. Shown are both the counts and normalized proportion of patients within each stratified ΔoncRNA level. **(C)** Number of oncRNA species detected in patient serum before (T0) and after (T3) neoadjuvant chemotherapy. **(D)** ΔoncRNA of patients grouped by clinically determined residual cancer burden (RCB) class. RCB 0 indicates pathological complete response while RCB III indicates high residual cancer burden. **(E)** Distant-metastasis free survival of patients grouped by ΔoncRNA. Also reported are the hazard ratio and *P* value based on a log-rank test. **(F)** Scatterplot of number of patients called as high ΔoncRNA versus resulting log-rank test $-\log_{10} P$ values using the cognate ΔoncRNA stratification. Points are colored by the resulting $\log_2$ hazard ratio. The ΔoncRNA threshold used for grouping high and low residual oncRNA burden in our reported survival analyses resulted in 27 high ΔoncRNA patients. **(G–H)** Forest plots of multivariate Cox proportional hazard model with ΔoncRNA and RCB class as covariates (G) and ΔoncRNA, subtype, and age as covariates (H). HER2 positive samples were excluded due to small sample size, and samples with missing clinical data were omitted.

#### Methods

##### Identification of oncRNAs in The Cancer Genome Atlas

11,082 TCGA small RNA-seq data were downloaded from the Genomic Data Commons in BAM format (hg38). Sample metadata was fetched using the GDC API. Reads were given a sequence complexity score using the DUST algorithm and removed from downstream analysis if the associated sequence complexity fell below a threshold (DUST score > 3) [42]. After conversion to BED format, unique small RNA loci across all samples were merged using mergeBed to create a comprehensive list of expressed small RNA loci. Loci longer than 200 base pairs were split via peak calling with SciPy (v.1.5), restricting loci peak lengths to be between 15 and 200 base pairs.

Non-cancerous extracellular and biofluid smRNA-seq data from the exRNA Atlas were downloaded in FASTQ format from the Gene Expression Omnibus (GEO) and the database of Genotypes and Phenotypes (dbGAP) and preprocessed in accordance with the cognate library preparation. Reads were then aligned to the genome (hg38) to generate BAM files. After applying the above low-complexity sequence filter, reads were converted to BED format. IntersectBed was used to create TCGA smRNA loci count tables for the exRNA Atlas samples. SmRNA loci observed in more than 7 exRNA Atlas samples were removed. The sample threshold was selected by using an elbow plot.

After filtering the TCGA smRNA loci by exRNA Atlas samples, we used the smRNA loci list to generate counts for each TCGA sample. The resulting smRNA loci counts, library size normalized counts (counts per million), and metadata for each sample were saved in a NoSQL database (MongoDB), aggregated and indexed by the smRNA loci.

To identify “orphan” smRNAs across TCGA, we first applied a filter to retain smRNAs that were largely absent in normal samples. Tumor-adjacent normal samples from TCGA were first stratified based on tissue type. SmRNAs that were observed in more than 10% of normal samples for any of the tissue types were removed. Only tissues with at least 10 normal samples were used for this normal tissue filtering step, which included 14 different tissue types. We then removed RNAs that were largely absent in cancer samples. For this step, we stratified cancer samples into 32 tissue types, and only retained smRNAs that were present in at least 10% of the cancer samples for at least one tissue type. For each cancer tissue type, we then used Fisher’s exact test to compare the presence and absence of the remaining smRNAs of tumor samples from the cognate cancer tissue type and normal samples from all tissue types. We selected smRNAs that were significantly present in the tumor samples of at least one tissue type, using an FDR cutoff of 0.1. After discovery of cancer-enriched smRNA loci, we then filtered our list of annotations against known smRNAs and miRNAs from publicly available annotations. SmRNAs overlapping by genomic coordinate with any of the existing annotations were removed. Lastly, we applied a filter using smRNA-seq libraries from 30 non-cancerous serum samples (cell-free RNA sequencing described below). Cancer-enriched smRNA loci that were detected in more than one of the samples were removed from our final annotated list of oncRNAs.

**Cancer tissue-of-origin modeling**

To evaluate the utility of oncRNA fingerprints for cancer tissue-of-origin modeling, we first split the TCGA samples into training and testing cohorts using a 80:20 train:test ratio, stratified by cancer types. We used the same methodology to train our classifier models on binarized, “digital” oncRNA profiles and normalized oncRNA expression profiles. Within the training cohort, we performed recursive feature elimination in a 5-fold cross validation scheme using a XGBoost classifier as our estimator to reduce the number of oncRNAs used as features from 260,968 to 1,805 (binary) features and 1,805 (cpm) features. After feature selection, we trained a final XGBoost classifier with 500 trees at max-depth of 3 on the full training cohort. The final model was evaluated on the held-out test set to calculate accuracy, precision, recall, and f1-scores.

**oncRNA and chromatin accessibility association analysis**

TCGA chromatin accessibility data were downloaded from GDC Publication Page (https://gdc.cancer.gov/about-data/publications/ATACseq-AWG). Of the 404 unique donors in the published study, 386 had matching TCGA smRNAseq data and were selected for inclusion in the analysis. Raw count matrices of published pan-cancer peaks of chromatin accessibility were normalized by library size. We then used intersectBed to identify ATAC peaks that overlapped with our set of oncRNA loci. To search for novel transcriptional activity, we removed any oncRNAs that overlapped with known genomic annotations, resulting in 10,725 oncRNA-ATAC peak pairs. For oncRNA-ATAC peak overlaps with at least 5 samples expressing the corresponding oncRNA, we performed an one-tailed Mann Whitney U test to test for higher ATAC peak scores in samples that expressed the cognate oncRNA compared to samples in which the oncRNA was not detected. *P* values were FDR corrected, resulting in 1,989 significant associations.

**Cancer subtype analysis and modeling**

Clinical metadata with subtype information for TCGA-BRCA datasets and TCGA-CRC (COAD and READ) were downloaded from cBioPortal(https://www.cbioportal.org/) and the Sage Bionetworks Synapse (https://www.synapse.org/), respectively. For each cancer, we used oncRNAs found to be statistically enriched in the cancer to train and evaluate XGBoost classifiers to predict cancer subtypes (Basal, Her2, Luminal A, and Luminal B for BRCA; CMS1, CMS2, CMS3, and CMS4 for CRC) in a 5-fold cross-validation setup. For both BRCA and CRC we used XGBoost classifiers with 100 trees at max-depth of 3. Performance metrics of the models including AUC of ROC, precision, recall, f1-score, and accuracy were averaged across folds.

**oncRNA selection for functional screens**

We triaged our list of ~260,000 of oncRNAs to select target oncRNAs for inclusion in our in-vivo over-expression and loss-of-function screens. oncRNAs were prioritized based on higher expression levels and prevalence across different cell line models of breast (MDA-MB231), colon (SW480), lung (A549), and prostate (C4-2B) cancers. Selected oncRNA loci longer than 38nt were trimmed to capture the region with the highest coverage or split into multiple smaller target loci if uniform coverage across the cell lines. The lengths of candidate oncRNA loci ranged from 15 to 38 nt after trimming for optimal performance in our TuD constructs.

**Library cloning**

For our combined oncTuD library, a library of 788 oligos (consisting of nominated oncRNAs as well as their corresponding TuD constructs) was designed and ordered from Twist Biosciences. The pool was resuspended to 5ng/uL final concentration in Tris-HCl 10mM pH 8, and a qPCR to determine Ct to be used for downstream library amplification was performed (forward primer: ATTTTGCCCCTGGTTCTT, reverse primer: CCCTAAGAAATGAACTGG) using a 16-fold library dilution.

**TuDs**

For TuDs, the library was then amplified via PCR and ran out on a 2% agarose gel to check library size (expected band of 200bp). PCR product was then cleaned up using a DNA Clean and Concentrator kit-5 (Zymo Research Cat. #D4003), and eluted in 25uL $H_2O$. Cleaned product was digested for 90 minutes using FD Esp3I (Thermo Fisher Cat. #FD0454). Digested inserts were run on a 8% TBE gel and extracted, and ethanol precipitated overnight in -20C. Inserts were then ligated into pUC6 (Addgene plasmid #49793) in a 100ng reaction with 1:1 insert:backbone ratio for 16hrs 16C. Ligated products were then ethanol precipitated overnight at -20C, and eluted in 4.5ul $H_2O$. 1.5ul ligation product was used for electroporation into 20ul MegaX DH10B $T1^R$ electrocompetent cells (Invitrogen Cat. #C640003), followed by maxiprep plasmid isolation.

5ug of intermediate pUC6 ligation product was then digested for 90 minutes using AgeI-HF (New England Biolabs Cat. #R3552S) and EcoRI-HF (New England Biolabs Cat. #R3101S). Digested inserts were then run on a 8% TBE gel, extracted, and then ethanol precipitated overnight at -20C. Inserts were then ligated into pLKO.1 (Addgene plasmid #10878) in a 100ng reaction with 1:1 insert:backbone ratio for 16 hrs at 16C. Ligated products were then ethanol precipitated overnight at -20C, and eluted in 4.5ul $H_2O$. 1.5ul ligation product was used for electroporation into 20ul MegaX DH10B $T1^R$ electrocompetent cells (Invitrogen Cat. #C640003), followed by maxiprep plasmid isolation.

**oncRNAs**

For oncRNAs, the library was then amplified via PCR and ran out on a 2% agarose gel to check library size (expected band of 75bp). PCR product was then cleaned up using a DNA Clean and Concentrator kit-5 (Zymo Research Cat. #D4003), and eluted in 25uL $H_2O$. Cleaned product was digested for 90 minutes using AgeI-HF (New England Biolabs Cat. #R3552S) and EcoRI-HF (New England Biolabs Cat. #R3101S). Digested inserts were ran on a 8% TBE gel and extracted, and ethanol precipitated overnight in -20C. Inserts were then ligated into pLKO.1 (Addgene plasmid #10878) in a 100ng reaction with 1:1 insert:backbone ratio for 16 hrs at 16C. Ligated products were then ethanol precipitated overnight at -20C, and eluted in 4.5ul $H_2O$. 1.5ul ligation product was used for electroporation into 20ul MegaX DH10B $T1^R$ electrocompetent cells (Invitrogen Cat. #C640003), followed by maxiprep plasmid isolation.

**Sequencing validation**

For sequencing validation, 300ng plasmid DNA was used as input to a first PCR targeting the oncTuD amplicon (forward primer: GGAAAGGACGAAACACCGGT; reverse primer: ATACTGCCATTTGTCTCGAGGTC) in 50ul volume, and PCR product was cleaned up using a Qiagen MinElute PCR purification kit, using a 1:1 volume of NTI cleanup buffer and eluting in 10ul volume (Qiagen Cat. #28004). 2ul of PCR product was then used as input into a second PCR to add Illumina adapter sequences (forward primer: ACACTCTTTCCCTACACGACGCTCTTCCGATCTGGAAAGGACGAAACACCGGT; reverse primer: GTGACTGGAGTTCAGACGTGTGCTCTTCCGATCTATACTGCCATTTGTCTCGAGGTC) in 50ul volume, and PCR product was cleaned up using Qiagen MinElute PCR purification kit with 1:4 NTI and eluting in 10ul volume. All 10ul of PCR product from the previous PCR was used as input into a final third indexing PCR to add Illumina indices (Illumina TruSeq UDI indices UDI009-0017). PCR product was cleaned up using 1X left-hand size selection (Zymo Cat. #D4084-4-10). Samples were then pooled and sequenced using a MiSeq v2 kit (Illumina Cat. #MS-102-2002).

**Lentivirus titration**

$2\times10^5$ cells per cell line (MDA-MB-231, SW480, C4-2B, A549) were seeded into 6-well plates (day 0). 24 hours post-seeding (day 1), 2 wells were counted and cell number per cell line recorded. To calculate titer, lentiviral library was added in an upwards range (100, 250, 500ul) in 3 wells per cell line. 72 hours post-seeding (day 3), puromycin was added to transduced wells, as well as an untransduced ‘kill’ well, at 8ug/mL final concentration. 3 days post-transduction (day 6), all wells were counted, as well as 2 untransduced and non-selected wells. Based on recorded cell number, one selected well per cell line (targeting 10-30% MOI) was used moving forward and expanded for future experiments.

**Cell preparation for subcutaneous injection**

Transduced cells were partitioned into 3 arms for our *in vivo* functional oncTuD screen. $2\times10^5$ cells per cell line were split into a 15cm plate for *in vitro* long-term passage, for purposes of growth normalization. $2\times10^5$ cells per cell line were also pelleted and frozen at -80C for downstream t0 gDNA extraction. For MDA cells, 16 million cells were resuspended to final concentration $1\times10^6$ cells/50ul in 1:1 PBS/matrigel, and bilateral mammary fat pad injections in 50ul final volume were performed in female, 8-12 week-old age-matched female NOD *scid* gamma (NSG) mice (n = 4). For SW480, C4-2B, and A549 cells, 16 million cells per cell line were resuspended to final concentration $1\times10^6$ cells/200ul in 1:4 PBS/matrigel, and bilateral subcutaneous injections in 200ul final volume were performed in either male (C4-2B) or female (SW480, A549) 8-12 week-old age-matched NSG mice (n =4 per cell line).

**Tumor gDNA extraction and library preparation**

3-4 weeks post-injection, tumors were harvested and processed using Quick-DNA midiprep plus kit (Zymo Research Cat. #D4075). For each processed tumor, gDNA was amplified in the ratio of 2.5ul input/25ul reaction volume in a first PCR targeting the oncTuD amplicon (forward primer: GGAAAGGACGAAACACCGGT; reverse primer: ATACTGCCATTTGTCTCGAGGTC). PCR product was cleaned up using 1X left-hand size selection (Zymo Cat. #D4084-4-10). 10% input from the first PCR was used in a second PCR to add Illumina adapter sequences (forward primer: ACACTCTTTCCCTACACGACGCTCTTCCGATCTGGAAAGGACGAAACACCGGT; reverse primer: GTGACTGGAGTTCAGACGTGTGCTCTTCCGATCTATACTGCCATTTGTCTCGAGGTC), and PCR product was cleaned up using 1X left-hand size selection (Zymo Cat. #D4084-4-10). 10% input from the second PCR was used in a last indexing PCR to add Illumina indices (Illumina TruSeq UDI indices UDI001-080), followed by 1X left-hand size selection (Zymo Cat. #D4084-4-10). Samples were pooled and sequenced on 2 lanes of NovaSeq SP200 150x8x8x50 at the UCSF Center for Advanced Technology (CAT).

**Cell culture**

All cells were cultured in a 37°C 5% CO2 humidified incubator. SW480 and C4-2B cell lines were cultured in RPMI-1640 medium supplemented with 10% FBS, glucose (2 g/L), L-glutamine (2 mM), 25 mM HEPES, penicillin (100 units/mL), streptomycin (100 µg/mL) and amphotericin B (1 µg/mL) (Gibco). MDA-MB-231 and A549 cell lines were cultured in Dulbecco's Modified Eagle Medium (DMEM) supplemented with 10% FBS, glucose (2 g/L), L-glutamine (2 mM), 25 mM HEPES, penicillin (100 units/mL), streptomycin (100 µg/mL) and amphotericin B (1 µg/mL) (Gibco). All cell lines were routinely screened for mycoplasma with a PCR-based assay**.**

**Target oncRNA expression and clinical association in TCGA-BRCA**

For oncRNAs with potential functional roles, we used the associated TCGA clinical metadata to compare their expression across tumor-adjacent normal tissue and cancer tissue and across breast cancer subtypes. We also stratified patients based on the expression levels of the oncRNAs and generated Kaplan-Meier curves. A log-rank test was used to compare the resulting survival curves.

**TCGA differential expression analysis and pathway analysis**

Raw gene expression data for the TCGA-BRCA dataset were downloaded from the Genomic Data Commons. Expression data were processed and normalized following the guidelines of the edgeR pipeline. Samples were grouped by presence or absence of cognate oncRNA and compared for differentially expressed genes using edgeR (v. 3.42.4), controlling for covariates including age and breast cancer subtype[43]. The resulting *P* values and log-fold change of each gene were used by iPage for pathway analysis to identify pathway perturbations associated with oncRNA expression[20].

**Orthotopic Tumor growth assay**

Tumor growth assays were performed by injecting cancer cells ($5\times10^5$ MDA-MB-231 or HCC1806 shctrl, oncRNA.ch7.29, or oncRNA.ch17.67) in 50µl 1:1 PBS:Matrigel (Corning) bilaterally into mammary fat pads of eight- to twelve-week old age-matched female NOD/SCID gamma mice. Tumor volume was assessed weekly by caliper measurements. Final tumor volume was measured *ex vivo* after surgically removing the tumor.

**Metastatic Lung Colonization Assay**

Eight- to twelve-week-old age-matched female NOD/SCID gamma mice (NSG, Jackson Labs, 005557) were used for lung colonization assays. For this assay, cancer cells constitutively expressing luciferase were suspended in 100 µL PBS and then injected via tail-vein ($1\times10^5$ MDA-MB-231 or HCC1806 shctrl, oncRNA.ch7.29, or oncRNA.ch17.67). Each cohort contained 4-5 mice, which in the NSG background is enough to observe a >2- fold difference with 90% confidence. Mice were randomly assigned into cohorts. Cancer cell growth was monitored in vivo at the indicated times by retro-orbital injection of 100 µl of 15 mg/mL luciferin (Perkin Elmer) dissolved in 1X PBS, and then measuring the resulting bioluminescence with an IVIS instrument and Living Image software (Perkin Elmer).

**Cell line mRNA sequencing and analysis**

mRNA-seq libraries were constructed using the QuantSeq 3' mRNA-Seq Library Prep Kit FWD according to the manufacturer's instructions (Lexogen, Cat. #015). RNA was extracted in replicates from MDA-MB-231 or HCC1806 shctrl, oncRNA.ch7.29, or oncRNA.ch17.67; 100-200ng RNA was then used as input to QuantSeq FWD. mRNA-seq libraries were pooled and sequenced on 1 lane of NovaSeqX 100x6x0x0 at the UCSF Center for Advanced Technology (CAT).

We then used cutadapt (v. 3.5) to remove adapter sequences. Preprocessed sequences were pseudoaligned to the transcriptome with Salmon (v. 0.14.1) to quantify gene expression. We used DESeq2 (v. 1.26.0) to perform the differential expression analysis with default settings [44]. *P* values were FDR corrected and used with gene expression data for pathway analysis with iPage, as mentioned above.

**Conditioned media collection and cell-free smRNA sequencing**

For each cancer of the 25 cancer lines, 200k-300k cells were seeded into a well of a 6-well plate in biological duplicate. After 48 hours, media was aspirated, cells were washed with PBS, then 3mL of fresh media prepared with exosome-depleted FBS was added. After 24 hours, conditioned media was collected, then cell-free RNA was extracted immediately with Quick-cfRNA Serum and Plasma kit (Zymo) and flash frozen. CfRNA was quantified with Qubit RNA HS, and ~14ng of each sample was used as input to construct small RNA-seq libraries with SMARTer smRNA-Seq Kit (Takara). For library prep, two modifications were made from the manufacturer's protocol: (a) the stock oligo dT for first strand synthesis was substituted for a custom primer with UMI's (5'CAAGCAGAAGACGGCATA CGAGATNNNNNNNNGTGACTGGAGTTCAGACGTGTGCTCTTCCGATCTTTTTTTTTTTTTTT-3') and (b) custom primers with single i5 indices were used for 18 cycles of cDNA amplification. For cleanup, the PCR products were column purified as per manufacturer's recommendations, and 175-300 bp PCR products were gel-purified from 8% polyacrylamide gels in TBE buffer. When necessary, the resulting libraries were additionally PCR-amplified with universal primers (5'-AATGATACGGCGACCACC-3' and 5'-CAAGCAGAAGACGGCATACGAG-3'). The libraries were sequenced on Illumina HiSeq 4000 or NovaSeq machines at the UCSF Center for Advanced Technology, on double-indexed single-end 50 nt runs.

We then used cutadapt (v1.15) to remove the poly(A) tails from the 3' end and 3 nucleotides unconditionally from the 5' end of each read to remove the template switch oligo. Reads with at least 15 base pairs after trimming were aligned to the human genome (hg38) using bowtie2 (v.2.3.5.1) with the end-to-end and sensitive setting. Libraries with UMIs were deduplicated using UMI-tools(v.1.1.0) with the default directional algorithm setting.The aligned BAM files were converted to BED format and intersectBed was used to quantify the number of reads mapping to known smRNAs (ie: miRNA, tRNA) and our list of annotated oncRNAs.

**I-SPY2 Trial and Clinical Samples**

All clinical blood samples were received from the I-SPY2 trial (NCT01042379), an ongoing, open-label, randomized, multicenter adaptive, phase 2 platform trial. Detailed description of the study design, patient eligibility and enrollment and oversight of the trial have been published previously[45,46]. The protocol for the I-SYP2 trial was approved by the Institutional Review Boards at all participating institutions. All patients signed written informed consent to participate in the trial and to allow the use of their biospecimens for research purposes.

Blood samples were collected at pretreatment (T0), and after NAC before surgery (T3) in marble/tiger-top vacutainer (serum separator) tubes. Tubes were placed upright for at least 15 minutes to properly clot. Within two hours of collection, tubes were centrifuged at 2500 rpm for 20 minutes at room temperature and then aliquoted into cryovial tubes and immediately frozen at -80C for storage.

**Serum RNA Extraction and sequencing**

For cell-free RNA extraction from patient serum samples, 0.5–1 mL of serum (stored at -80C from collection to extraction) per sample was used. The samples were thawed at room temperature and RNA was extracted using Quick-cfRNA Serum and Plasma kit (Zymo) following manufacturer's recommendations, eluted in 15 µl nuclease-free water and stored at -80C. Small RNA-seq libraries were constructed, sequenced and analyzed as described above for cell line conditioned media cell-free RNA.

**ISPY-2 survival analysis**

Residual oncRNA burden (ΔoncRNA) for each patient was calculated as:

$$\Delta\text{oncRNA} = N_{T3} - N_{T0}$$

where $N_{T0}$ and $N_{T3}$ were the total number of oncRNA species detected per million reads sequenced from the serum samples at time point 0 (prior to neoadjuvant chemotherapy) and time point 3 (completion of neoadjuvant chemotherapy treatment and prior to surgery), respectively. Patients were stratified by ΔoncRNA levels into two groups: i) high and persistent residual oncRNA burden and ii) low residual oncRNA burden (Fig. S4F). Using these stratifications we generated Kaplein-Meier curves and performed a log-rank test to calculate the associated *P* value. We used multivariable Cox regression analysis to assess ΔoncRNA as an independent predictor of survival after NAC while controlling for established clinical variables. To account for the sample size, we performed several iterations of Cox analysis with different covariates separately: ΔoncRNA with pCR, ΔoncRNA with RCB class, and ΔoncRNA with age and breast cancer subtype.
